# Cryo-EM structures of *Cereibacter sphaeroides* ferrochelatase reveal an oligomeric state that occludes the substrate-binding site

**DOI:** 10.64898/2026.11.02.762883

**Authors:** Jack H. Salisbury, Pu Qian, Yibo Jin, Amanda A. Brindley, Joshua R. Barrett, Nathan B. P. Adams, Ruby R. E. Harvey, Frederick R. Hawkings, Harold A. Cannon, George A. Sutherland, Matthew P. Johnson, James D. Reid, David J. K. Swainsbury, Pablo Castro-Hartmann, Per A. Bullough, C. Neil Hunter, Andrew Hitchcock

**Author notes:** Authors for correspondence (JHS), (CNH) & (AH).

## Abstract

Protoporphyrin IX (PPIX) is the last shared intermediate of heme and (bacterio)chlorophyll ((B)Chl) biosynthesis. Insertion of iron into PPIX by the PPIX ferrochelatase (FeCh) PpfC produces heme, whereas magnesium insertion by magnesium chelatase (MgCh) initiates (B)Chl synthesis. Chlorophototrophic organisms must regulate this biosynthetic node to ensure the correct flux of PPIX to each of these crucial tetrapyrrole products, presenting a particularly acute problem for the facultative bacterium *Cereibacter* (previously *Rhodobacter*) *sphaeroides*, which can switch between chemotrophic (respiratory) and phototrophic (photosynthetic) growth modes. To elucidate the structural basis for the catalytic and regulatory functions of *C. sphaeroides* FeCh (PpfC*^Cs^*), we used cryogenic electron microscopy to determine its structure, to our knowledge the first such information for a Gram-negative or phototrophic organism. We determined the 2.6 Å structure of an active site mutant of PpfC*^Cs^*, which forms a dimer with a monomer-to-monomer interface that coordinates two hemes. We also determined a 2.5 Å structure of the wild-type enzyme, in which two dimers interact in a homotetrameric arrangement that occludes PPIX binding at the active site of each monomer. We propose that the PpfC*^Cs^* dimer likely docks onto the inner membrane in a manner analogous to the human enzyme, whereas the tetramer represents a transiently inactive state whereby the hydrophobic lips and active site are both occluded. These novel oligomerization states of FeCh may have evolved to influence the flux of tetrapyrroles through the heme/BChl branchpoint in purple bacteria.

## Introduction

Chlorophototrophic organisms require a diverse range of tetrapyrroles to support their growth, including hemes, (bacterio)chlorophylls ((B)Chls) and cobalamin (vitamin B_12_). Each of these macrocyclic, metalated compounds is derived from the universal precursor 5-aminolevulinic acid (ALA), which is synthesized from succinyl-CoA and glycine in the C4/Shemin pathway or from glutamate in the C5 pathway (1). ALA, which is specifically utilized for tetrapyrrole biosynthesis, is sequentially converted to the last intermediate common to the synthesis of all tetrapyrroles, uroporphyrinogen III (UroII). In the classical heme biosynthesis pathway in eukaryotes and Gram-negative bacteria, UroIII is decarboxylated to generate coproporphyrinogen III and then protoporphyrinogen IX, the latter of which is then oxidised to form protoporphyrin IX (PPIX); iron insertion into PPIX produces protoheme (heme B) whereas chelation with magnesium generates Mg-PPIX in the first dedicated step of (B)Chl biosynthesis. While it was long considered that this PPIX route was the only way to make heme, alternative siroheme-dependent (2) or coproporphyrin III-dependent (3) pathways are present in Gram-positive bacteria and Archaea.

The metal chelatase enzymes that insert divalent cations into their respective macrocycles are essential for tetrapyrrole biosynthesis and are broadly classified as either type I or type II (4). Type I chelatases are heteromeric, ATP-dependent enzymes that insert magnesium (Mg) during (B)Chl biosynthesis or cobalt (Co) in the aerobic (or late Co-insertion) vitamin B_12_ biosynthesis pathway (5, 6). Conversely, type II chelatases are generally single-subunit monomeric or homodimeric enzymes that do not require ATP for metal ion insertion (4). There are several type II chelatases: PPIX ferrochelatase (FeCh) (PpfC, previously HemH); coproporphyrin III FeCh (CpfC), which inserts iron into coproporphyrin III in the coproporphyrin III-dependent heme biosynthesis pathway; SirB, which inserts iron into sirohydrochlorin (SHC) in the final step of siroheme biosynthesis; CbiK and CbiX, which insert cobalt into SHC in the anaerobic (or early insertion) vitamin B_12_ biosynthesis pathway; and CfbA, which catalyzes nickel insertion into SHC during coenzyme F_430_ biosynthesis (1). In addition to their central catalytic roles, metal chelatases are often found at nodes in the tetrapyrrole biosynthesis pathway; for example, PpfC and magnesium chelatase (MgCh) compete for PPIX, and SirB, CbiX/K and CfbA all act on SHC (1). It is therefore critical that these branchpoints are regulated to ensure the correct allocation of tetrapyrrole intermediates to each downstream pathway. For example, in chlorophototrophic organisms the assembly of reaction centre and light-harvesting complexes requires large amounts of (B)Chls but heme is also an essential cofactor for both photosynthetic and respiratory electron transport chains, as well as many other important enzymes, thus PPIX must support both (B)Chl and heme biosynthesis.

While it is well known that heme negatively regulates tetrapyrrole biosynthesis by inhibiting ALA synthase at the beginning of the pathway (7, 8), regulation of the heme/(B)Chl branchpoint is less well understood. *In vitro* enzyme assays reveal higher rates of iron insertion into PPIX by PpfC than magnesium insertion by MgCh (9, 10), which does not align with the higher proportion of (B)Chl required for phototrophic growth. This problem is particularly acute for purple phototrophic bacteria (PPB) such as *Cereibacter* (previously *Rhodobacter*) *sphaeroides*, which can transition between chemotrophic (respiratory) and phototrophic (photosynthetic) modes of growth. Considering there is no evidence for the downregulation of FeCh at the gene or protein level during the switch to phototrophic growth in *C. sphaeroides* (11, 12), and PPB lack the MgCh-activating Gun4 protein found in oxygenic phototrophs (13), one possibility is that PpfC activity is regulated post-translationally. Consistent with this hypothesis, PpfC*^Cs^* forms a protein complex with the small membrane protein PufQ (14). The *pufQ* gene is part of the photosynthesis gene cluster, specifically the *pufQKBALMX* operon that encodes components of the reaction center-light-harvesting 1 (RC-LH1) core complex (15), suggesting a direct link between photosynthetic gene expression and FeCh regulation. Deletion of *pufQ* resulted in a decrease in BChl levels under low-oxygen growth conditions, accompanied by excretion of the coproporphyrinogen III precursor from the cell (14). However, BChl synthesis and photoheterotrophic growth were recovered by a missense suppressor mutation in the *ppfC* gene, which was found to markedly decrease FeCh activity *in vitro*, consistent with PufQ playing a role in regulation of the heme/BChl branchpoint.

Although there are structures of the PpfC enzymes from humans (*Homo sapiens*, PpfC*^Hs^*: **Fig. S1**) and yeast (*Saccharomyces cerevisiae*, PpfC*^Sc^*) (16, 17), and CpfC from the Gram-positive bacterium *Bacillus subtilis* (18), there is no such information for Gram-negative bacteria. Yet, in the case of *C. sphaeroides*, the PpfC structure would provide a framework for understanding the regulation of the heme/BChl branchpoint. Accordingly, we determined electron cryomicroscopy (cryo-EM) structures of the *C. sphaeroides* FeCh (PpfC*^Cs^*), which forms a homodimer with an interface that coordinates two heme molecules using a His/Cys pair of axial ligands to the central iron. Two of these homodimers can self-associate to form a tetramer with residues that extend into the active site cleft, appearing to prevent substrate binding. The structure of a PpfC*^Cs^*variant with increased affinity for PPIX shows how substrate binding to the active site prevents tetramer formation, allowing the active site to interface with the inner membrane to receive substrate from the preceding enzyme in the pathway, the membrane-integral protoporphyrinogen IX dehydrogenase (PPO). We discuss the possible implications of these novel oligomerization states of FeCh relating to the flux of PPIX to both Bchl and heme biosynthesis during phototrophic growth.

## Results

### Structure of dimeric *Cereibacter sphaeroides* ferrochelatase

Recombinant N-terminally-His-tagged PpfC*^Cs^* (UniProt accession: Q3IYK5) was purified following production in *Escherichia coli*. We used a PpfC*^Cs^* E320K variant, equivalent to the inactive human E343K PpfC mutant that has enhanced affinity for PPIX (19) and purified the enzyme in the presence of sodium cholate as was reported previously for other FeChs (see methods) **(Fig. 1*A*)**. The absorbance spectra of isolated PpfC*^Cs^*E320K showed distinct peaks at 360 and 414 nm consistent with oxidized heme, with additional features at 507 and 546 nm, characteristic of PPIX **(Fig. 1*B*)**, although the PPIX spectral features are less pronounced than those observed previously for other FeChs (e.g. <u>20</u>). The spectra of PpfC*^Cs^*further contrasts with that of the human enzyme, which has a C-terminal extension containing a cysteine-rich motif (CxxCxxxxC) that coordinates a [2Fe-2S] cluster and gives rise to characteristic absorbance features of PpfC*^Hs^* at 330, 460 and 550 nm (21) **(Fig. S1)**.

**Figure 1:**
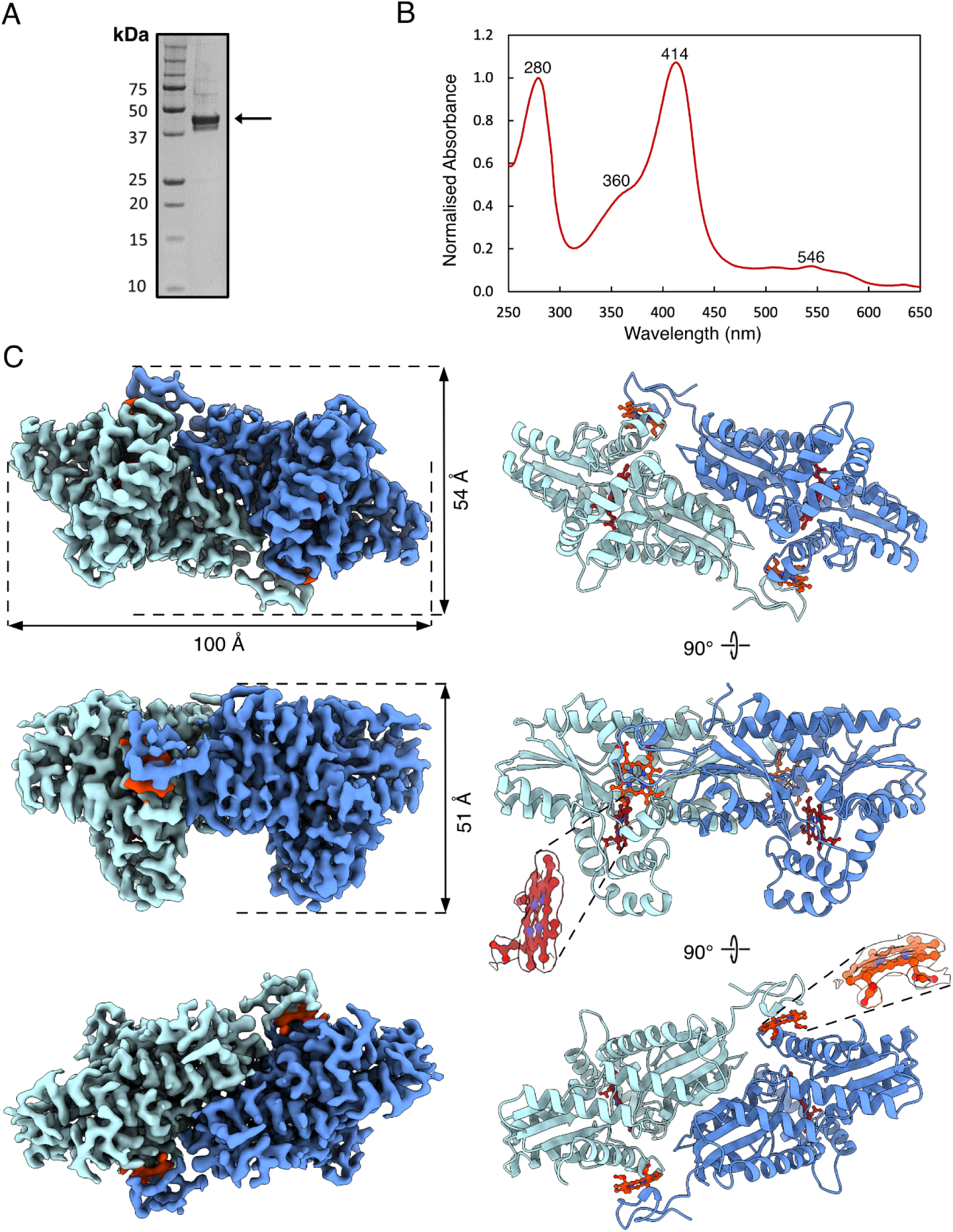
Structure of the E320K variant of *C. sphaeroides* ferrochelatase. (*A-B*) Coomassie stained SDS-PAGE gel **(*A*)** and absorption spectrum **(*B*)** of purified E320K PpfC*^Cs^*. The predicted molecular weight of His-tagged E320K PpfC*^Cs^* is 46 kDa, indicated by the black arrow in panel (A). **(*C*)** Refined C2 map and corresponding model of E320K PpfC*^Cs^*. Individual subunits are shown in shades of blue, with bound hemes colored in dark orange (interface) and red (active site).

Purified E320K PpfC*^Cs^* was used to prepare grids for structure determination by cryo-EM. Due to issues with preferential orientation, movies from 11,370 un-tilted grids and 5,456 tilted (35°) grids were collected. Following motion correction, 5.4 million particles were selected for iterative two-dimensional (2D) and three-dimensional (3D) classifications and a single high-resolution class containing 96,153 particles was refined to 2.6 Å resolution with C2 symmetry imposed **(Fig. S2-S3)**. An AlphaFold3 (22) model predicted from two PpfC units was then docked into the final B-factor sharpened map and iteratively refined in PHENIX **(Table S1)**. Cryo-EM density for Met1 to Glu19 at the N-terminus and the preceding His-tag was absent, presumably due to high flexibility of this region. The resulting structure of the E320K PpfC*^Cs^* variant revealed a novel dimer interface for PpfC **(Fig. 1*C*),** with each PpfC subunit rotated 180° relative to its partner. The interface between subunits was found to contain density for two metalated porphyrins, which we model as hemes, consistent with the absorbance spectrum of the purified enzyme. The active site, contained within the enzyme’s three lip regions, was occupied by a second porphyrin species, which was also shown to contain clear density for a chelated metal ion within the macrocycle **(Fig. S4)**. Thus, our structurally determined E320K mutant appears to represent a post-turnover, product-bound state of PpfC.

### Comparison of the active site in PpfC*^Cs^* and PpfC*^Hs^*

The structure of each PpfC*^Cs^* monomer displays conserved architecture with PpfC*^Hs^* (Root Mean Square Deviation (RMSD) = 3.4 Å across all 324 pairs) **(Fig. S5)**, with an active site cleft flanked by symmetric α4/β4 domains (16). This cleft is protected by a hydrophobic ‘cap’ formed by the upper and lower lip regions (UL and LL, respectively) **(Fig.2*A*)**. In PpfC*^Cs^* the UL hairpin region is formed by α helices 1-3 (residues 58-117). A lower lip region (residues 240-252, LL_1_) encompasses the short α8 helix and is directly adjacent to His239, equivalent to the catalytically important His263 residue in PpfC*^Hs^* (23), and a second lower lip loop region (LL_2_) is formed from residues 280-291. These three lip regions act to tightly coordinate the porphyrin within a hydrophobic cavity with two small solvent exposed cavities to the right side of the macrocycle **(Fig.2*B*)**. This inter-locked conformation also appears to occlude the porphyrin from external solvent, with a hydrogen bond network observed between Tyr77 (UL), Arg244 (LL_1_) and Glu285 (LL_2_) **(Fig.2*C*)**. This interlocked lip region additionally allows hydrophobic residues on the UL and LL_2_ regions (Trp82/Leu85/Phe282) to form a barrier between the bound porphyrin and the external solvent/membrane. The propionate-6 group of the bound porphyrin is coordinated by Arg94 (UL) and the substituted Lys320 residue, while the propionate-7 group is coordinated by Tyr102 (UL) and Thr52 (β1/α1 loop) **(Fig.2*D**-E*)**. The additional coordination of the propionate-6 group by Lys320 explains the increased affinity of the E320K variant for porphyrin, as reported previously for the E343K variant of the human enzyme (19). Deprotonation of the PPIX macrocycle prior to metalation has been proposed to occur through a His/Glu pair in the human enzyme (23); and we highlight a pair of glutamates (Glu324/Glu328) adjacent to the equivalent His239 and Glu320 residues in PpfC*^Cs^*that may provide a pathway for proton release to the external solvent **(Fig. S6)**. Active site residues around the porphyrin were generally well conserved with the human variant **(Fig.2*F*)**, except for Met76 in PpfC*^Hs^*, which is Leu50 in PpfC*^Cs^*. In PpfC*^Hs^* Met76 is suggested to participate in iron insertion in the human enzyme (24, 25), and the implications of its absence from PpfC*^Cs^* are discussed later.

**Figure 2:**
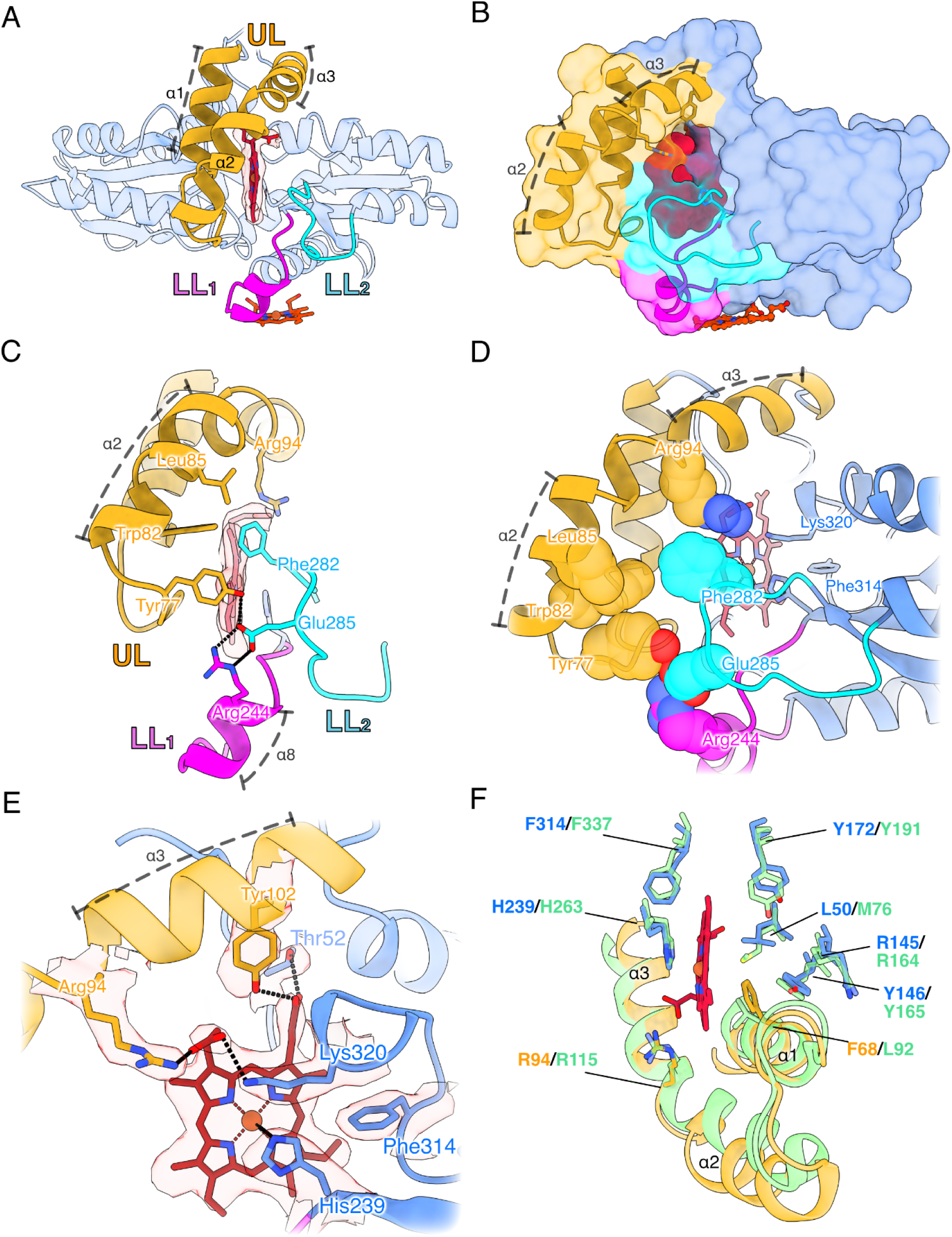
Comparison of the active site of *C. sphaeroides* PpfC with the human enzyme. (*A*) Topology of a monomer of PpfC*^Cs^* highlighting the lip regions that participate in porphyrin binding (in this case heme, shown in red with the central iron atom in orange); upper lip (UL, gold), lower lip region 1 (LL_1_, magenta), and lower lip region 2 (LL_2_, cyan). **(*B*)** Surface representation of PpfC*^Cs^*, showing how the lip regions result in a hydrophobic cleft to accommodate porphyrin binding, colored as in (A). The heme bound at the dimer interface is shown in dark red (see Figure 3D-E for more details). **(*C*)** Side view of the porphyrin binding region. Phe282 (LL_2_), Trp82 (UL) and Leu85 (UL) form a hydrophobic ‘cap’ to the active site cleft. The lip regions interact through a hydrogen bonding network between Arg244 (LL_1_), Glu285 (LL_2_) and Tyr77 (UL). **(*D*)** Second view of the hydrophobic cap formed by the three lip regions, with residues shown as spheres. **(*E*)** Side view of the bound porphyrin, showing coordination of the central metal by His239. The propionate-6 group is coordinated by Arg94 (UL) and Lys320, while the propionate-7 group is coordinated by Tyr102 (UL) and Thr52. **(*F*)** Superimposition of the PpfC*^Hs^* E343K substrate-bound ‘closed’ structure (PDB: 2HRE, green) and PpfC*^Cs^* (colored as in panels A-E). The active site porphyrin from the PpfC*^Cs^* E320K structure is shown for reference (red).

### A novel dimer interface in PpfC*^Cs^* coordinates hemes

The dimer interface of PpfC*^Cs^* is formed via hydrophobic interactions between adjacent β4 strands in each monomer, such that an extended β-sheet stabilizes the dimer **(Fig. 3*A-B*)**. This architecture is possible because of the lack of the C-terminal extension seen in eukaryotic FeChs **(Fig. S1)**, which would otherwise prevent two monomers associating in this way **(Fig. S7)**. Similarly, the extended N-terminus of PpfC*^Cs^*prevents interaction of the α helices (α9 in PpfC*^Cs^*/α11 in PpfC*^Hs^*) that form the dimer interface in the human enzyme **(Fig. S7)**. A total of 34 putative hydrogen bonds form the PpfC*^Cs^* dimer interface; of note are hydrogen bond networks between Arg201/Asn364 at the external face of the dimer **(Fig. 3*B*)**, as well as Gln197/Asp185, and Arg196/Glu205, which form symmetrical salt bridge networks either side of the β4 strands **(Fig. 3*C*)**. We observed additional density for a heme at the dimer interface, bound within a pocket proximal to the α9 helix **(Fig. 3*D*)**. This heme is tightly bound in a cleft formed between the two monomers, consisting of several residues on the N-terminal loop of one monomer (Arg27, His30, His35, Leu38) and the α8/α9 helices of the adjacent monomer (Tyr207, Lys243, Leu246, Met247, His253, Cys254, Lys258, Arg261). The central metal of the heme is coordinated by Cys254 on the α9 helix and His35 on the extended N-terminal region of the adjacent monomer **(Fig. 3*D*)**; individual substitution of these residues to Ser or Ala, respectively, abolished heme binding **(Fig. S8)**. The 6-propionate group of the heme is hydrogen bonded by Arg27 (on the N terminal loop), Lys258 (on the α9 helix) and Tyr207 (sits on the β4:α7 loop), while the 7-propionate group is coordinated solely by Arg261 on the α9 helix **(Fig. 3*E*)**.

**Figure 3:**
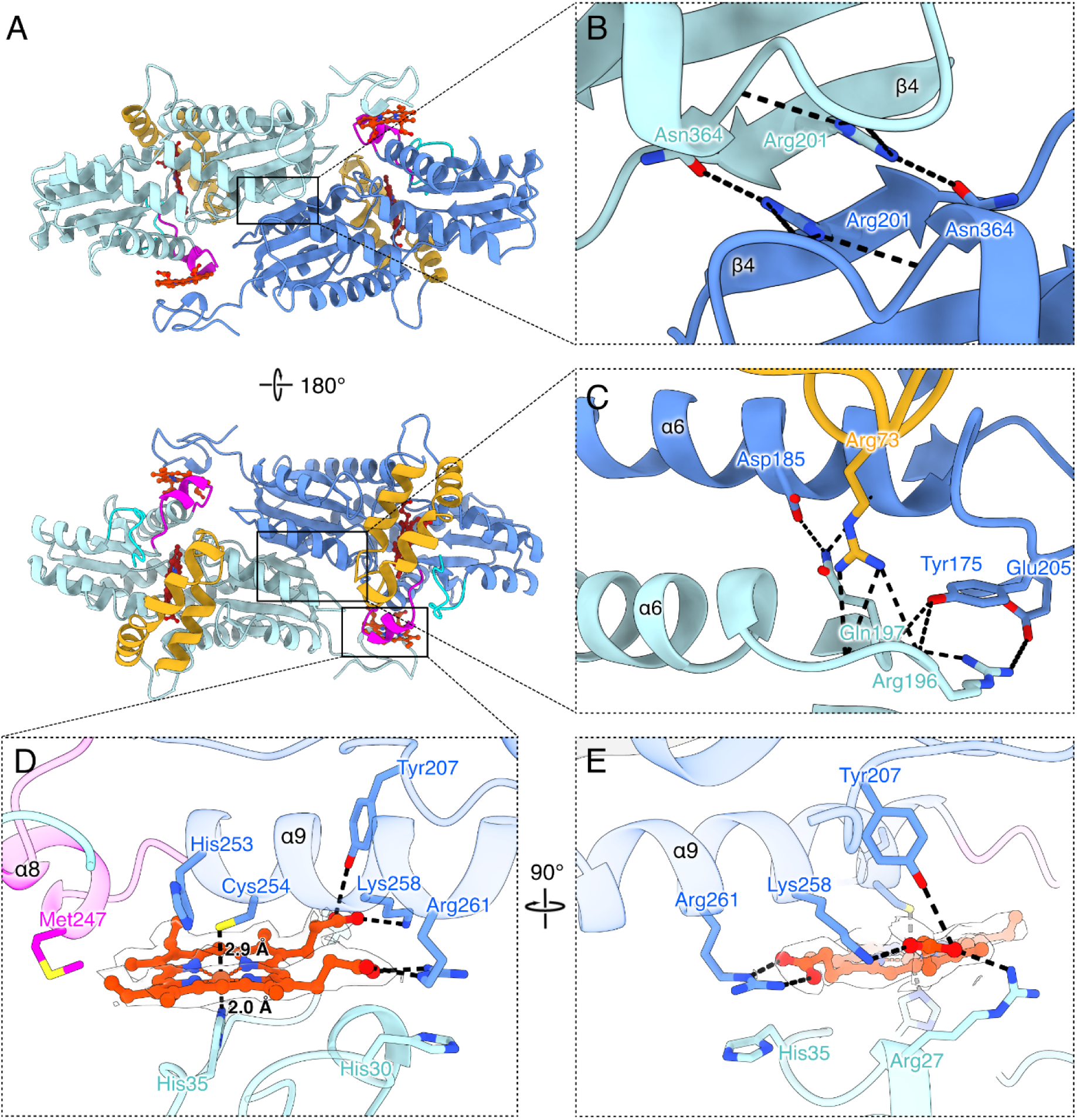
The dimer interface of PpfC*^Cs^.* (*A*) Cartoon representation of the distinct dimer interface of E320K PpfC*^Cs^*. **(*B*)** Hydrogen bond networks between the β4 strands, consisting of interactions between Asn364 and Arg201. **(*C*)** Hydrogen bonding network centred around the α6 helix of each subunit, consisting of interactions between Asp185/Gln197 and Glu205/Arg196. Additionally, Arg73 of the UL is shown to form hydrogen bonds to the backbone of the α6/β4 loop of the adjacent subunit. **(*D-E*)** Heme coordination at the dimer interface. Cys254 and the His35 from opposing monomers act as axial ligands to the heme iron atom, while Arg27, Tyr207, Lys258, and Arg261 form ligands to the propionate groups.

### The wild-type PpfC*^Cs^* enzyme forms a homotetramer

In contrast to the E320K variant, a significant proportion of the wild-type (WT) *C. sphaeroides* enzyme was present in the soluble fraction following production in *E. coli*. To understand the biochemical basis for this increase in solubility, we purified wild-type PpfC*^Cs^* (as for E320K but in the absence of detergent) **(Fig. 4*A*)**. The absorbance spectra of the purified wild-type enzyme showed distinct peaks at 360 and 420 nm, compared with peaks at 360 and 414 nm for E320K PpfC, and lacked features corresponding to PPIX **(Fig. 4*B*).** This WT preparation was shown to be active in a spectroscopic assay measuring the insertion of ferrous iron into PPIX **(Fig. 4*C*)**.

**Figure 4:**
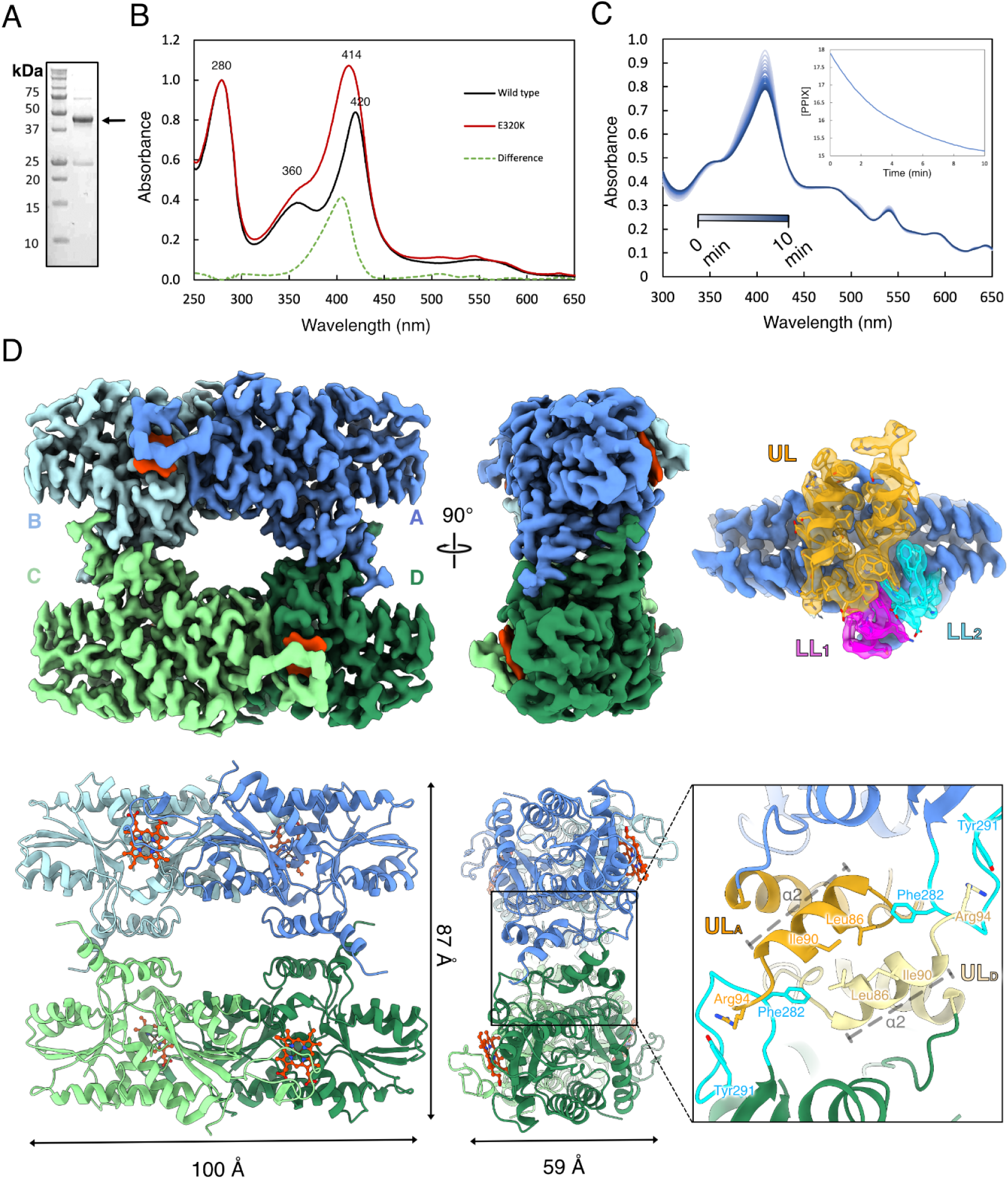
Structure of wild-type *C. sphaeroides* ferrochelatase. (*A-B*) Coomassie stained SDS-PAGE gel **(*A*)** and absorption spectrum **(*B*)** of wild-type PpfC*^Cs^*. In ***(B)*** the E320K PpfC*^Cs^* and the E320K minus wild type difference spectra are also shown. The predicted molecular weight of His-tagged PpfC*^Cs^* is 44 kDa, indicated by the black arrow in panel (A). **(*C*)** PPIX metalation by purified PpfC*^Cs^* monitored by the decrease in absorbance of the Soret band at 407 nm. With 20 μM PPIX the enzyme had a specific activity of 50.23 ± 1.29 μmol min^-1^ mg^-1^. **(*D*)** Density map for PpfC*^Cs^*, with opposing dimers shown in shades of blue (A/B) and green (C/D), with the interface hemes shown in dark orange. The lower lip regions are colored magenta (LL_1_) and cyan (LL_2_) and the upper lip region (UL) in gold. **(Inset)** Representation of the α2 helices of opposing monomers, which constitute the primary hydrophobic inter-dimer interface of the PpfC*^Cs^* tetramer. The interface is buttressed by the Phe282 residue on the LL_2_ region of each subunit. Arg94, at the terminal end of the α2 helix, is oriented towards the LL_2_ region of the opposing monomer, forming a putative hydrogen bond with Tyr291.

To determine the structure of wild-type PpfC*^Cs^*, we collected 3,337 movies of untilted samples **(Table S1)** and picked 1,893,803 particles from the processed, motion-corrected micrographs, followed by a series of iterative iterative 2D- and 3D-classifications. The particles appeared ring-shaped and symmetric in the initial classes, so D2 (dihedral) symmetry was imposed on the 3D classification, yielding a single, high-resolution class containing 89,047 particles, which was refined to a global resolution of 2.5 Å **(Fig. S9-S10)**. The segmented cryo-EM density map revealed PpfC*^Cs^* formed a dimer-of-dimers, with opposing dimer pairs interacting primarily via the hydrophobic lip regions proximal to the active site **(Fig. 4*D*)**. No density corresponding to PPIX was observed at the active site, although densities for the two hemes found at the interface within each homodimeric pair in the E320K structure were present. Each subunit within the homotetrameric structure adopts a similar fold to the monomers in the E320K dimer **(Fig. 5A)**, except for the UL region, with the density for residues Phe96-Met116 that form the α3 helix and part of the α4 helix missing in the structure of the wild-type enzyme **(Fig. S11)**.

**Figure 5:**
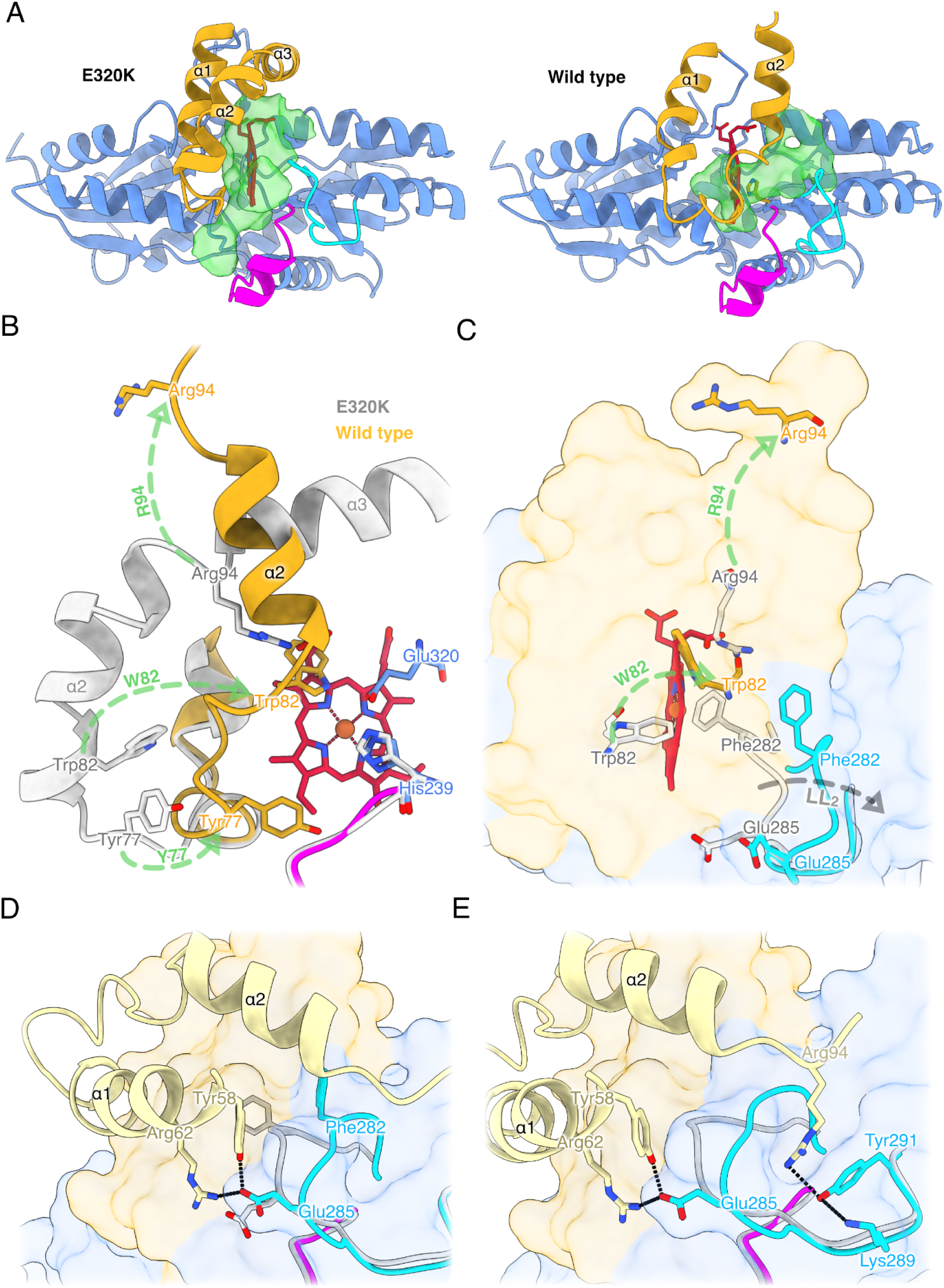
Comparison of the wild-type and E320K PpfC*^Cs^* structures reveals how the upper lip controls PPIX entry to the active site. (*A*) Cartoon representation of E320K and wild-type PpfC*^Cs^*, with lip regions colored as in Figure 2. The active site cleft of each subunit is shown in green and the position of the E320K porphyrin is also shown in the wild-type model for reference. The wild-type PpfC*^Cs^* structure shows the clear change in conformation of the upper lip, with the effect of decreasing the size of the active site cleft such that porphyrin binding is no longer possible. **(*B*)** Superimposition of the UL from the wild-type (gold) and E320K (grey with crimson porphyrin) PpfC models. To form the tetramer, the α2 helix rotates inwards towards the active site cleft. This conformational change causes Trp82 and Tyr77 to orient inwards, occluding the porphyrin-binding site. Secondly, Arg94, which coordinates propionate-6 of the bound porphyrin in the E320K model, moves approximately 20 Å. **(*C*)** Rotated view of the superimposed structures, showing movement of Trp82 and Arg94. The LL_2_ loop also swings outwards, changing the relative positions of Phe282 and Glu285. **(*D*)** Surface view of a monomer from the wild-type PpfC structure with the LL_2_ loop highlighted. The movement of this loop allows for the formation of a hydrogen bond network between Glu285 and residues from the UL of the opposing subunit (Tyr58 and Arg62, pale gold). **(*E*)** Arg94 of the opposing subunit’s UL region forms a hydrogen bond network with Tyr291 and Lys289 of the LL_2_ region.

The UL and LL_2_ regions mediate the interaction between two PpfC*^Cs^* dimer pairs within the tetramer; viewing the inter-dimer interface from a second symmetry axis, it is apparent that the α2 helices of each opposing monomer form the primary interaction surface via a hydrophobic leucine-rich patch that is further stabilized by Phe282 of the LL_2_ region **(Fig. 4D, inset)**. The identification of this higher oligomeric state provides a clear rationale for the increase in solubility of the wild-type enzyme, with the membrane-interfacing hydrophobic UL region now forming an inter-dimer interaction.

### The active site is occluded in the tetrameric structure

While the overall topology of a PpfC monomer was similar in the wild-type and E320K variants (RMSD of 0.52 Å excluding lip regions), the conformation of the UL region was distorted in the wild-type enzyme such that the α2 helix is oriented away from the active site cleft and towards the opposing dimer **(Fig. 5*A*)**. This UL movement has a significant effect on the active site cavity, which is reduced in volume in the wild-type enzyme **(Fig. 5*A*)** such that PPIX binding is sterically blocked. Closer analysis of the UL region revealed that the α2 helix had collapsed inwards and was tilted in the opposite direction relative to its position in the structure of the E320K variant **(Fig. 5*B*)**. This UL conformational change results in Trp82 facing inwards towards the active site, just 3.0 Å from Glu320, effectively occluding the substrate binding site **(Fig. 5*B-C*).** The UL rearrangement additionally causes a dramatic shift in the position of the propionate-coordinating Arg94 residue, which moves approximately 20 Å away from the position it adopts in the active site of E320K PpfC*^Cs^* **(Fig. 5*B*-*C*)**. The LL_2_ lip region also undergoes a conformational change in wild-type PpfC*^Cs^* (RMSD of 3.5 Å between residues 280-291), moving outwards relative to the active site cleft **(Fig. 5*C*),** allowing Glu285 to participate in the inter-dimer interface by hydrogen bonding to Tyr58 and Arg62 of the UL of the opposing monomer **(Fig. 5*D*)**. Lys289 and Tyr291 of LL_2_ participate in a hydrogen bond network with the displaced Arg94 of the opposing monomer, providing stability to the active site-occluded structure **(Fig. 5*E*)**.

## Discussion

### PpfC*^Cs^* adopts substrate-free tetrameric and substrate-bound dimeric forms

PpfC is a crucial enzyme in the biosynthesis of heme B, which is a versatile cofactor ubiquitous across nearly all organisms on Earth. This work represents, to our knowledge, the first structural study of this enzyme from a photosynthetic organism, in addition to the first report of a PPIX FeCh structure from a Gram-negative bacterium. The topology of the PpfC*^Cs^* monomer shows good agreement with previous studies on the human and yeast enzymes (16, 17), with a conserved bilobal architecture that is thought to have evolved from a single lobed ancestor protein (26, 27). However, we demonstrate that this bacterial variant oligomerizes in two novel ways. Firstly, PpfC*^Cs^* dimerizes via a stacked β-sheet motif, supported by a N-terminal extension that contributes to the interface by coordination of a heme molecule **(Fig. 3)**. This dimerization contrasts with PpfC*^Hs^*, where dimer pairs are associated via α11 helices (α9 in PpfC*^Cs^*), but still allows the active site lips of each subunit to orient in the same direction. We suggest that this dimerisation allows for both PpfC*^Cs^* units to interact with the membrane surface as is proposed for eukaryotic PpfC (28), explaining the need for sodium cholate in the purification of both of these variants. Structural alignments reveal how the respective extensions of each enzyme (the N-terminal extension of PpfC*^Cs^* and the C-terminal extension of PpfC*^Hs^*) have resulted in the convergent evolution of separate dimeric states both retaining the capability to interface with the membrane surface **(Fig. S7)**.

Purification of wild-type PpfC*^Cs^* in the absence of detergent enabled the identification of a second oligomeric state, in which two dimer pairs interact via the upper and lower lip regions, forming a tetramer with D2 symmetry **(Fig. 4*D*)**. As these regions form the main hydrophobic interface with the inner membrane (29), we propose that this interaction disengages the enzyme from the membrane, enabling detergent-free isolation of the soluble tetrameric complex. We show that formation of this second inter-dimer interface requires a significant conformational change in both the UL and LL_2_ regions, including the disordering of the UL ɑ3 helix **(Fig. 4*D***, **Fig. 5*A*)**. The changes that allow for the formation of this interface have the secondary effect of orientating hydrophobic residues deep within the active site cavity, effectively blocking access of the PPIX substrate **(Fig. 5*B*)**. We suggest that both the unique dimer interface and absence of C terminal extension of PpfC*^Cs^* are key determinants in the formation of the tetramer, with the alternate PpfC*^Hs^* dimer sterically prevented from adopting a similar oligomeric state.

A distinct feature of Ppf*^Cs^* is the presence of bound heme at the dimer interface, with the central metal coordinated by two axial ligands: an external cysteine on the α9 helix (Cys254) of one monomer, and a histidine residue on the N terminal loop (His35) of the other. A bis-axial Cys/His pair is unusual for heme binding but has been observed in the *b*-type heme of cystathionine β synthase (30) and the *c*-type hemes of SoxAX cytochromes (31). In the former enzyme, it has been shown that the absence of heme results in a decrease in activity due to mis-folding and aggregation (32). As we were unable to assay the H35A and C254S heme binding mutants of Ppf*^Cs^* due to decreased solubility and aggregation, it is reasonable to suggest the heme cofactor plays a similar role in enzyme stability in the ferrochelatase, and the presence of these residues may infer the ability of purple bacterial PpfC homologues to form the dimer. An alignment of purple bacterial PpfC sequences **(Fig. S12)** showed that the axial coordinating residues for the heme are conserved in *Rhodobacter capsulatus*, *Dinoroseobacter shibae*, *Roseobacter denitrificans*, *Rhodovulum sulfidophilum* and *Rubrivivax gelatinosus*, but are absent in both *Rhodopseudomonas palustris* and *Rhodospirillum rubrum*. However, we could not find conservation for these residues across a range of non-photosynthetic Gram-negative bacteria **(Fig. S13)**, suggesting heme binding is a novel adaptation of some purple bacterial FeChs Interestingly, CbiK^P^, a periplasmic cobaltochelatase (CoCh) homologue from the sulfate-reducing bacterium *Desulfovibrio vulgaris*, is a tetrameric heme-binding protein, but binds hemes in the central cavity of the tetramer using a pair of histidine residues not conserved in other CoChs (33). The cellular localization of this protein suggests it is involved in iron metabolism/heme transport rather than tetrapyrrole biosynthesis (34).

### The PpfC*^Cs^* active site is well conserved and appears to stably bind product in the E320K enzyme

The active site of PpfC*^Cs^* shows good general agreement with that of the PPIX FeChs from yeast and humans **(Fig. S5)**, with good spatial conservation of the invariant His/Glu pair proximal to the macrocycle (proposed to carry out the deprotonation event) and a phenylalanine (Phe314 in PpfC*^Cs^*) at the anterior of the cleft, which has been suggested to act as a molecular ‘anvil’ for the initial distortion of the porphyrin (24). We found less conservation on the opposite side of the macrocycle ring, with the human Met76 residue substituted by Leu50 in PpfC*^Cs^*. A M76A mutation was found to severely lower activity in PpfC*^Hs^*(24), with this methionine proposed to carry out weak iron coordination preceding metalation. While it is unlikely that Leu50 plays a role in metalation, we did find strong conservation for this residue in both purple bacterial enzymes **(Fig. S12)** and more widely PpfC from Gram-negative bacteria **(Fig. S13)**. Comparing the active site porphyrin of E320K PpfC*^Cs^* to the analogous E343K PpfC*^Hs^*variant (19), we observed density within the porphyrin macrocycle of E320K PpfC*^Cs^* that was not observed in the human structure. The authors demonstrated that the human E343K was inactive; whereas the *C. sphaeroides* E320K mutant appears to be capable of undergoing a single turnover event based on our structure. The π helix (residues 340-360 in PpfC*^Hs^*), a region that has been shown to unwind to facilitate product release (35, 36), is in the wound conformation in both WT and E320K PpfC*^Cs^* **(Fig. S5)**.

### The role of PpfC in regulation of the heme/bacteriochlorophyll branchpoint in tetrapyrrole metabolism

Chlorophototrophs face the challenge of allocating metabolic flux between the heme and (B)Chl pathways, particularly since fluctuating light levels and other environmental factors alter the demand for (B)Chls. This is particularly acute in facultative bacteria such as *C. sphaeroides*, which must also regulate this heme/BChl branchpoint when they transition between their major chemotrophic (respiratory) and phototrophic (photosynthetic) growth modes in response to changes to oxygen tension and illumination (37). Under microoxic conditions that pertain to phototrophic growth, *C. sphaeroides* dramatically increased BChl production (via magnesium metalation of PPIX by MgCh) as well as doubling the amount of cellular heme (14). Our current study suggests that this organism may be able to influence the ratio of heme/Bchl via oligomeric switching of its PpfC from membrane-accessible to soluble states. PpfC regulation has been observed in oxygenic phototrophs, in which FeCH possesses a conserved C-terminal Chl-binding motif (CAB domain) (38, 39). Chl binding to this domain of cyanobacterial PpfC leads to its dimerisation, whereas the pigment-free enzyme is monomeric (40, 41, 42). Despite these different mechanisms, it is clear that PpfC activity must be tightly regulated in order to optimize (B)Chl production in photosynthetic organisms.

An earlier study of regulation of the heme/BChl branchpoint uncovered a role for the membrane intrinsic peptide PufQ and a membrane protein of unknown function, Rsp_6067, which form a complex with PpfC*^Cs^* (14). Chidgey *et al.* showed that BChl synthesis was severely decreased in a Δ*pufQ* mutant but was restored by a suppressor mutation in PpfC*^Cs^* (Y58H), which significantly lowered FeCh activity in *in vitro* assays. This led to the suggestion that PufQ promotes BChl biosynthesis by somehow limiting the participation of FeCh at the heme/BChl branchpoint to enhance activity of the slower MgCh enzyme. Our present work shows that Tyr58 of PpfC*^Cs^*contributes to the PpfC*^Cs^* inter-dimer interface **(Fig. 5*D*)**. Therefore, one possibility is that the lower activity of the Y58H mutant is due to increased stability of the tetrameric form, which would require a conformational change to interface with the inner membrane to access PPIX, thereby favoring the synthesis of BChls. However, further native biochemical analyses to assess the dimer/tetramer ratio of the enzyme in the presence and absence of substrate is required to validate this assumption.

### Experimental Procedures Purification of PpfC*^Cs^*

To purify recombinant *C. sphaeroides* FeCh, the *ppfC* gene was amplified from the *C. sphaeroides* 2.4.1 genome by PCR (see **Table S2** for primer sequences) using Q5® High-Fidelity DNA Polymerase (NEB). The gene was ligated into the NdeI and XhoI sites of the pET-28a(+) vector (Novagen) creating the plasmid pET-28a::*ppfC^C^*^s^, and the gene sequence was verified by automated Sanger sequencing (Eurofins). The pET-28a::*ppfC^C^*^s^ construct was introduced to *E. coli* BL21(DE3) and 1 L LB cultures in 2 L baffled conical flasks containing 50 µg/mL kanamycin were grown at 37 °C and 250 rpm to an optical density at 600 nm (OD_600_) of ∼0.6, at which point *ppfC* expression was induced by the addition of isopropyl β-d-1-thiogalactopyranoside (IPTG) to a final concentration of 0.4 mM. The cultures were cooled to 18 °C and incubated for a further 16 h at 250 rpm prior to cell harvesting by centrifugation (4,000 x*g*, 4 °C, 20 min). The pellets were resuspended in ∼ 30 mL binding buffer (50 mM HEPES pH 8, 200 mM NaCl, 5 mM imidazole) per L of original culture, and a cOmplete EDTA-free protease inhibitor tablet (Roche Holding AG) and a few crystals of DNAse I (Merck) were added. Cells were broken by probe-sonication (12 rounds of 30 s on, 30 s off, 70% amplitude) in an ice water bath using a SONICS vibra-cell, before isolation of the soluble cell-free extract (CFE) as the supernatant following centrifugation (48,000 x*g*, 4 °C, 20 min).

N-terminally His_6_-tagged PpfC*^Cs^* was purified using a combination of immobilised metal affinity chromatography (IMAC), ion exchange chromatography (IEC) and size exclusion chromatography (SEC). The CFE was first applied to 5 mL of pre-equilibrated Ni-NTA resin (Cytivia) and sequentially washed with 20 column volumes (CVs) of binding buffer with 20 mM or 40 mM imidazole, respectively, prior to elution in 2 CVs of the same buffer containing 200 mM imidazole. The red-colored elution fraction was concentrated to a volume of 2.5 mL in a Vivaspin®20, 30 kDa MWCO centrifugal filter (Cytiva) and exchanged into buffer A (50 mM HEPES pH 8) using a PD-10 desalting column (Cytiva). The sample was diluted to a volume of 25 mL in the same buffer and PpfC*^Cs^* was further purified by IEC using a 5-mL HiTrap Q HP column (Sigma) run at 2 mL min^-1^ with a gradient of 0-100% Buffer B (50 mM HEPES pH 8, 1 M NaCl) over 20 CVs collecting 1 mL fractions. Red-colored fractions were analyzed by SDS-PAGE and those enriched in PpfC*^Rs^* (typically with an A420/A280 ratio >1.5) were pooled and concentrated to a volume of 1 mL in a Vivapsin®20, 30 kDa MWCO centrifugal filter (Cytiva) prior to a final SEC purification step using a HiLoad 16/600 Superdex column (Sigma) pre-equilibrated and run in 50 mM HEPES pH 8, 200 mM NaCl at 0.5 mL min^-1^ collecting 2 mL fractions. Red-colored fractions containing PpfC*^Cs^* were analyzed by SDS-PAGE, pooled and concentrated as above. For long-term storage 20% (w/v) glycerol was added and 50 µL fractions of protein at a concentration of ∼ 500 µM were flash frozen in liquid nitrogen and stored at −80°C. Protein concentration was determined using the Expasy-predicted extinction coefficient (https://web.expasy.org/protparam/) of the His-tagged PpfC*^Cs^* enzyme of 65,320 M^-1^ cm^-1^.

Plasmids for production of H35A and C254S substituted variants of PpfC*^Cs^* were generated using the QuikChange II site-directed mutagenesis kit (Agilent) with the pET-28a::*ppfC^C^*^s^ plasmid as template and purification was attempted as for the wild-type enzyme (see **Table S2** for primer sequences).

### Purification of PpfC*^Cs^* E320K

A construct to produce the E320K variant of PpfC*^Cs^* was initially made in the pET-28a::*ppfC^C^*^s^ plasmid using the QuikChange II site-directed mutagenesis kit (Agilent) but attempted purification using the same conditions for wild-type PpfC*^Cs^* described above did not yield soluble His-tagged protein, so we switched to the method used previously for the human enzyme (16, 19). The *ppfC*^E320K^ gene was amplified and ligated into the BamHI and HindIII sites of the pTrcHisA vector (provided by Professor Amy Medlock, University of Georgia) such that the produced protein would have an extended N-terminus including a His_6_-tag. The sequence-verified construct was introduced to *E. coli* BL21(DE3) and 1 L Terrific Broth (20 g/L tryptone, 24 g/L yeast extract, 0.4% (w/v) glycerol, 0.17 M KH_2_PO_4_, 0.72 M K_2_HPO_4_) cultures in 2 L baffled conical flasks supplemented with 100 µg/mL ampicillin were grown overnight at 30 °C and 180 rpm, with expression of *ppfC* driven from the leaky *trc* promoter in the absence of IPTG induction. Cells were harvested by centrifugation (4,000 x*g*, 4 °C, 20 min) and the pellets were resuspended in binding buffer (as above but with the addition of 1% (w/v) sodium cholate). PpfC E320K was initially purified by IMAC as described above but with 1% (w/v) sodium cholate added to the wash buffers. The IMAC elution was further purified by IEC and SEC as described above, but IEC buffers A and B contained 1% (w/v) sodium cholate, and the SEC buffer contained 0.1% (w/v) sodium cholate. Protein concentration was determined using the Expasy-predicted extinction coefficient (https://web.expasy.org/protparam/) of the His-tagged PpfC*^Cs^* E320K enzyme of 72,310 M^-1^ cm^-1^.

### Purification of PpfC*^Hs^*

The human FeCh (PpfC*^Hs^*) was purified for spectral comparison with PpfC*^Cs^* (see **Fig. S1C** for the absorbance spectrum of PpfC*^Hs^*). A gene encoding PpfC*^Hs^*lacking its 54-residue N-terminal transit peptide (https://www.uniprot.org/uniprotkb/P22830/entry) was synthesized as a gBlock by Integrated DNA Technologies and cloned into the NheI and HindIII sites of pTrcHisA such that the recombinantly produced protein contained an N-terminal His_6_-tag to aid purification. The protein was purified in the presence of sodium cholate as described for the E320K variant of PpfC*^Cs^*.

### Ultraviolet-Visible (UV/Vis) spectroscopy

UV-Vis absorption spectra were measured on a Cary 60 spectrophotometer (Agilent) with a scan rate of 600 nm s^-1^.

### Ferrochelatase activity measurements

Ferrochelatase activity was assayed using the insertion of ferrous (Fe(II)) iron into protoporphyrin IX (PPIX) (Sigma). Assays were performed in a total volume of 400 µL in micro-cuvettes (BRAND®), containing 0.2 μM PpfC, 20 μM PPIX and 10 μM FeCl_2_ in an assay buffer consisting of 25 mM Tris pH 7.4 with 0.15% (v/v) Tween 80. PPIX was prepared by dissolving 3 mg of PPIX (Sigma) in 40 mM NaOH, 1 % (v/v) Tween 80, 100 mM MOPS pH 7, 40 mM HCl. Each buffer component was added stepwise and mixed by vortexing for 30 s. The final mixture was centrifuged at 14,000 xg for 30 minutes at 4 °C and the supernatant was taken and stored. The stock concentration was determined in 2.6 M HCl using the extinction coefficient ε = 297, 000 M^-1^ cm^-1^ at 408 nm (43). All reagents other than PpfC were combined and equilibrated at 30 °C prior to the addition of enzyme to initiate the reaction, which was monitored using either the *Scanning Kinetics* program on a Cary60 spectrophotometer scanning 300-650 nm at 3 scans min^-1^ for 25 minutes or by monitoring single wavelength absorbance at 407 nm to measure the decrease in the Soret band of PPIX as it becomes metalated. An endpoint reading at 612 nm was taken and the concentration of PPIX over time was calculated as ((A407 - A612) / ε) using the calculated assay extinction coefficient for PPIX, which we determined as ε = 41.5 mM^-1^ cm^-1^ at 407 nm in assay buffer.

### Cryo-EM grid preparation

A Vitrobot Mark IV (ThermoFisher) was used for freezing grids. Protein solution (4 µl at a concentration of 250 μM) was applied to a glow discharged 300 mesh Quantifoil R1.2/1.3 Cu grid (Quantifoil Micro Tools GmbH). Grids were glow discharged using a GloQube (Quorum Technologies) with a current of 50 mA for 60 sec. Grids were incubated in the sample chamber at 4 °C with 99% relative humidity and blotted for 3 seconds with a blot force of 3 and then plunge frozen in liquid ethane before moving to liquid nitrogen. Grids were stored under liquid nitrogen prior to screening and data collection.

### Cryo-EM data collection

Both datasets were collected on a Titan Krios G3i (Thermo Fisher Scientific) equipped with a Falcon 4i direct electron detector and Selectris X energy filter (Thermo Fisher Scientific). The microscope was operated at 300 kV with an energy slit width of 15 eV and a nominal magnification of 165,000 (0.734 Å/pix for wild-type PpfC*^Cs^*and 0.729 Å/pix for E320K PpfC*^Cs^*). For wild-type PpfC*^Cs^*, a total of 3,337 movies were collected with a total dose of 40 e*^-^*/Å*^2^* under a defocus value range from −0.6 to −2.0 μm. For E320K PpfC*^Cs^*, a total of 11,370 movies were collected at 0° tilt, and a further 5,456 movies were recorded at a tilt of 35°, with a total dose of 60 e*^-^*/Å*^2^* under a defocus value range from −0.6 to −2.0 μm.

### Cryo-EM data processing

Wild-type PpfC*^Cs^* was processed using Relion 4.1 (44). Beam induced movement was corrected by MotionCor2 (45) with 5 x 5 patch, dose weighting and a B factor of 150. All default parameters in CTFFIND 4.1.14 (46) were used for CTF correction. 1,893,803 particles were initially picked up using Topaz 0.2.5 (47) with a model trained on approximately 500 particles. Particles were extracted using a box size of 288 pixels (downscaled to 96 pixels, binning = 3) and subjected to multiple rounds of reference-free 2D classification (T-factor = 2, Class =200). Selected classes that appeared as ring-like particles with an approximate diameter of 10 × 8 × 3 nm. The 457,885 selected particles were used to generate a *de novo* initial model. This model was used as a reference for unsupervised 3D classification (T = 4, 4 classes), with the resulting classes appearing to have strong D2 symmetry. Thus, a single class was aligned along its D2 symmetry axis and used as a reference for a second round of 3D classification (T = 4, 8 classes) with D2 symmetry imposed. Particles from one class that showed clear secondary structural features were re-extracted in a box size of 288 pixels (unbinned) and subjected to multiple rounds of 3D refinement with two rounds each of per-particle CTF refinement (fitting magnification, defocus, astigmatism, B-factor, phase-shift, beam tilt, trefoil and fourth order aberrations). A final Bayesian polishing step was applied before a final 3D refinement to achieve an estimated resolution of 2.51 Å based on the gold standard Fourier Shell Correlation (FSC) at 0.143 (48).

E320K PpfC*^Cs^* was processed in CryoSPARC 4.5.3 (49). Beam-induced motion was corrected using patch motion correction in CryoSPARC live and CTF parameters of motion corrected micrographs were estimated using patch CTF, also in cryoSPARC live. Approximately 5,400,000 particles were picked using the automated blob-picker tool (70-100 Å, NCC of 0.35+, power rating 350-1050) and extracted using a box size of 280 pixels (downscaled to 70 pixels, binning = 4). These particles were subjected to multiple rounds of 2D classification, selecting separately for particles representing smaller ‘end’ views and larger ‘side’ views. In total, 212,736 particles representing possible side views and 275,257 particles representing possible end views were combined and subjected to *ab initio* reconstruction and non-uniform refinement, followed by heterogeneous refinement with 2 classes. The particles from the best resolved class produced an anisotropic map due to preferential orientation. To obtain missing particle views, the micrographs were re-picked using circular blobs (100-160 Å, NCC of 0.25+, power rating 700-1181). Particles were extracted using a box size of 400 pixels (downscaled to 100 pixels, binning = 4) and subjected to multiple rounds of 2D classification. Only views not present in the previous particle sets were kept, with 95,138 particles of the new side view kept and combined with the previously good particle sets. Duplicated particles were removed using the remove duplicates job in cryoSPARC. 171,563 particles were re-extracted (unbinned), with a non-uniform refine job giving a consensus map with a resolution of 2.9 Å. This particle set was subjected to one round of 3D classification using 2 classes and filtered resolution of 6 Å. The best class containing 96,153 particles was used for non-uniform refinement with C2 symmetry applied to give an estimated final resolution of 2.63 Å based on the gold standard FSC at 0.143 (48).

### Model building and refinement

The initial model of PpfC*^Cs^* was predicted using ModelAngelo (50) with four copies of the PpfC protein sequence as the input. This model was course-fit into the D2 map for model building and refinement. *De novo* building of the disordered N-terminus was done in *Coot* (51), with successive refinements done in *Coot* and PHENIX (52). Coordinates for the heme were downloaded from the SMILES database and were docked into the map within *Coot.* The initial model of the E320K dimer was predicted using AlphaFold3 (22), using two copies of the PpfC protein as the input. This model was docked into the C2 refined density map and refined iteratively in *Coot* and PHENIX. A total of 108 waters were modelled into the wild-type PpfC*^Cs^* map.

## Data availability

The EM maps have been deposited to the Electron Microscopy Data Bank (EMDB, https://www.ebi.ac.uk/emdb/) under the accession codes EMD-57912 (PpfC*^Cs^*) and EMD-58452 (PpfC*^Cs^* E320K) and the protein models to the RCSB protein data bank (PDB, https://www.rcsb.org/) under the accession codes 30PK (PpfC*^Cs^*) and 31JV (PpfC*^Cs^* E320K).

## Acknowledgements

AH acknowledges the support of a Royal Society University Research Fellowship (award references URF\R1\191548 and URF\R\241006), which also funded JHS and AAB. JRB acknowledges a White Rose Doctoral Training Partnership PhD studentship (project reference 2881069) funded by the Biotechnology and Biological Sciences Research Council (BBSRC). FRH was supported by a Diamond Light Source PhD studentship (award number STU0355) jointly funded by the University of Sheffield. PQ, AAB, GAS and CNH acknowledge support from the European Research Council (Synergy Award 854126). DJKS acknowledges start-up funding from the University of East Anglia. We additionally thank the School of Biosciences Cryo-Electron Microscopy Facility at the University of Sheffield for access for grid screening and computational support.

## Notes

### Competing Interest Statement

The authors have declared no competing interest.

