## Supplementary Tables S1-S2 and Supplementary Figures S1-S13 for "Cryo-EM structures of *Cereibacter sphaeroides* ferrochelatase reveal an oligomeric state that occludes the substrate-binding site"

**Supporting information (Supplementary Tables S1-S2 and Supplementary Figures S1-S13)**

Jack H. Salisbury[1,2]*, Pu Qian[1,3], Yibo Jin[3], Amanda A. Brindley[1], Joshua R. Barrett[1], Nathan B. P. Adams[1,4], Ruby R. E. Harvey[1], Frederick R. Hawkings[1,5,6], Harold A. Cannon[1,7], George A. Sutherland[1,8], Matthew P. Johnson[1], James D. Reid[9], David J. K. Swainsbury[10], Pablo Castro-Hartmann[3], Per A. Bullough[1], C. Neil Hunter[1,]* and Andrew Hitchcock[1,]*

[1] School of Biosciences, University of Sheffield, Sheffield S10 2TN, UK

[2] Current affiliation: Medical Research Council Mitochondrial Biology Unit, University of Cambridge, Cambridge CB2 0XY, UK

[3] Materials and Structural Analysis, Thermo Fisher Scientific, 5651 GG Eindhoven, Netherlands

[4] Current affiliation: NanoTemper Technologies, 81369 Munich, Germany

[5] Diamond Light Source Ltd, Oxfordshire OX11 0DE, UK

[6] Current affiliation: Medical Research Council Laboratory of Molecular Biology, Cambridge CB2 0QH, UK

[7] Current affiliation: Department of Chemistry, University of Oxford, Oxford OX1 3TA, UK

[8] Current affiliation: Baseimmune, London BioScience Innovation Centre (LBIC), London NW1 0NH, UK

[9] School of Mathematical and Physical Sciences, University of Sheffield, Sheffield S3 7HF, UK

[10] School of Biological Sciences, University of East Anglia, Norwich NR4 7TJ, UK

**Table S1. Cryo-EM data collection, refinement, and validation statistics.**

| | PpfC | E320K |
|---|---|---|
| | PDB-30PK; EMD-57912 | PDB-31JV; EMD-58452 |
| **Data collection and Processing** | | |
| Magnification | 165,000 X | 165,000 X |
| Electron exposure (e-/Å$^2$) | 40 | 60 |
| Defocus range | -0.6 to -2.0 µm | -0.6 to -2.0 µm |
| Pixel size (Å) | 0.734 | 0.729 |
| Symmetry imposed | D2 | C2 |
| Initial particle images (no.) | 1.8 million | 5.4 million |
| Final particle images (no.) | 89,047 | 96,153 |
| Map resolution (Å) | 2.51 | 2.63 |
| FSC threshold | 0.143 | 0.143 |
| Model resolution range (Å) | 1.74 to 3.24 | 2.38 to 3.90 |
| Map sharpening B factor (Å$^2$) | 64.5 | 71.6 |
| **Model Composition** | | |
| Non-hydrogen atoms | 21,676 | 11,670 |
| Protein residues | 1328 | 706 |
| Waters | 108 | – |
| Ligands | HEM: 4 | HEM: 4 |
| Bond lengths RMSD (Å) | 0.007 | 0.007 |
| Bond angles RMSD (°) | 1.484 | 1.010 |
| **Validation** | | |
| Model-map CC | 0.8177 | 0.7896 |
| MolProbity score | 1.02 | 1.25 |
| Clash score | 2.16 | 4.00 |
| Rotamer outliers (%) | 0.00 | 0.00 |
| **Ramachandran plot** | | |
| Favoured (%) | 97.87 | 97.72 |
| Allowed (%) | 2.13 | 2.28 |
| Outliers (%) | 0.00 | 0.00 |

**Table S2. List of primers used in this study.** Restriction sites used for cloning are underlined.

| Name | Sequence (5′→3′) |
|---|---|
| PpfC$^{Cs}$_28a_F | CCCCATATGACGCCGATGCTGCAAGCG (NdeI) |
| PpfC$^{Cs}$_28a_R | GCGCCTCGAGGCCGCGCGAGCCAG (XhoI) |
| PpfC$^{Cs}$_pTrcHis_F | AGTCGGATCCACGCCGATGCTGCAAGC (BamHI) |
| PpfC$^{Cs}$_pTrcHis_R | AGTCAAGCTTCTTCGGGCCTCTCTCAGC (HindIII) |
| PpfC$^{Cs}$_H35A_F | GGCAGCTCCGGAGCGTCTGCGGGCGC |
| PpfC$^{Cs}$_H35A_R | GCGCCCGCAGACGCTCCGGAGCTGCC |
| PpfC$^{Cs}$_C254S_F | CGCACTGGCTATGGTAGGGATCGCCC |
| PpfC$^{Cs}$_C254S_R | GGGCGATCCCTACCATAGCCAGTGCG |
| PpfC$^{Cs}$_E320K_F | CTCTTCCAGCGTCTTGATGCAGTCCGCCG |
| PpfC$^{Cs}$_E320K_R | CGGCGGACTGCATCAAGACGCTGGAAGAG |

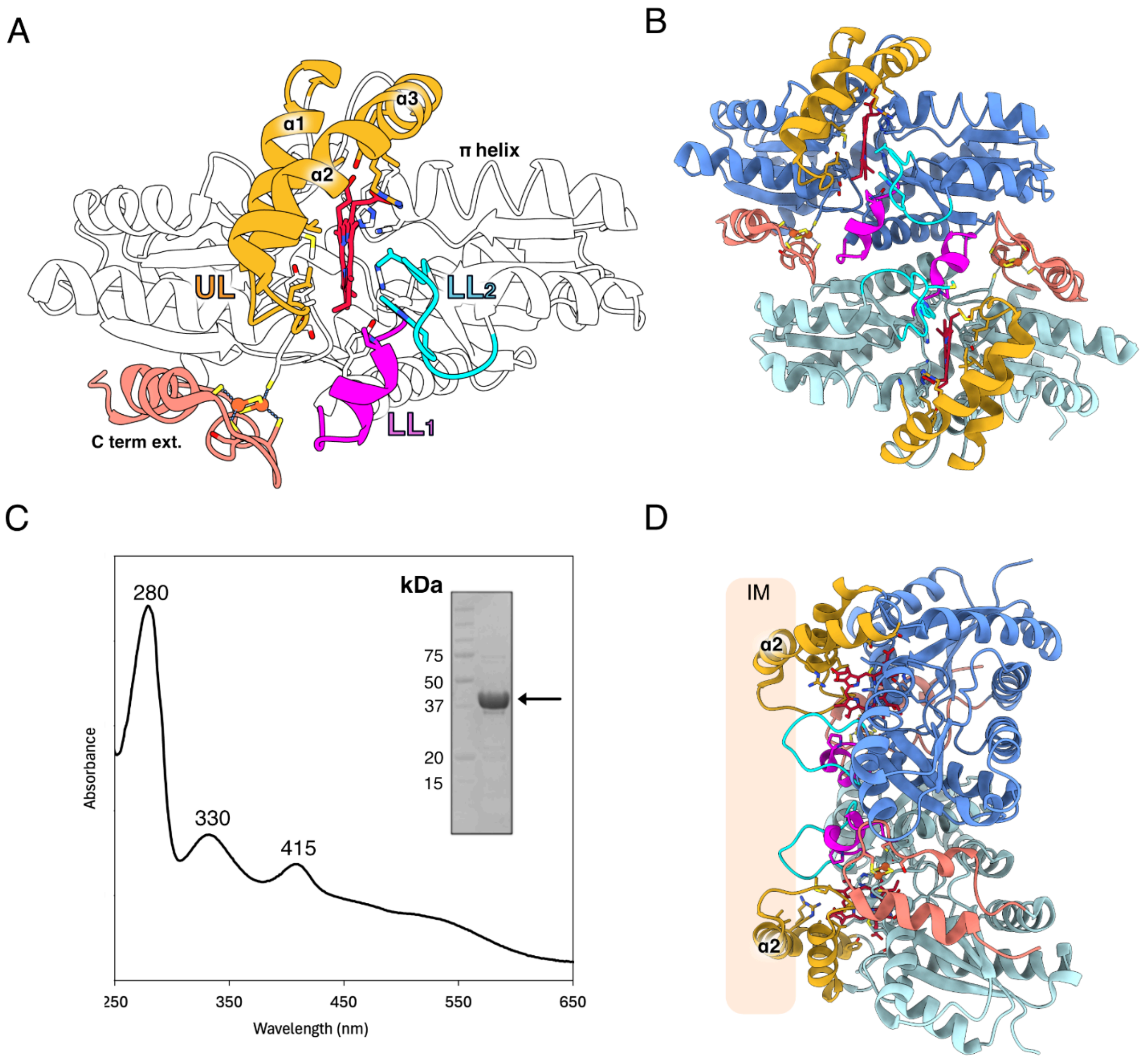


**Figure S1: Topology of the *Homo sapiens* PpfC enzyme**. (***A***) Cartoon representation of a monomer of the substrate-bound E343K variant of PpfC$^{Hs}$ (PDB: 2HRE) showing the symmetric alpha/beta domains that flank the active site cleft. The three lip regions (UL = upper lip, $LL_1$ = lower lip loop 1, $LL_2$ = lower lip loop 2) are shown in gold, magenta, and cyan, respectively. The extended C terminus (salmon) that coordinates the [2Fe-2S] cluster (orange-yellow) is labelled, as well as the π helix, which has been suggested to 'unwind' to facilitate product release (35, 36). (***B***) Cartoon representation of human E343K mutant, indicating the dimer interface mediated by the the α11 helices and C terminal regions. (***C***) Absorbance spectrum of wild-type PpfC$^{Hs}$ purified in this study. The feature at 330 nm is due to the [2Fe-2S] cluster, and that at 415 nm is assigned to heme bound at the active site. (***D***) Rotated view of the human dimer interface, showing how the hydrophobic upper lip regions (gold) facilitate the association of the enzyme with the inner membrane (IM).

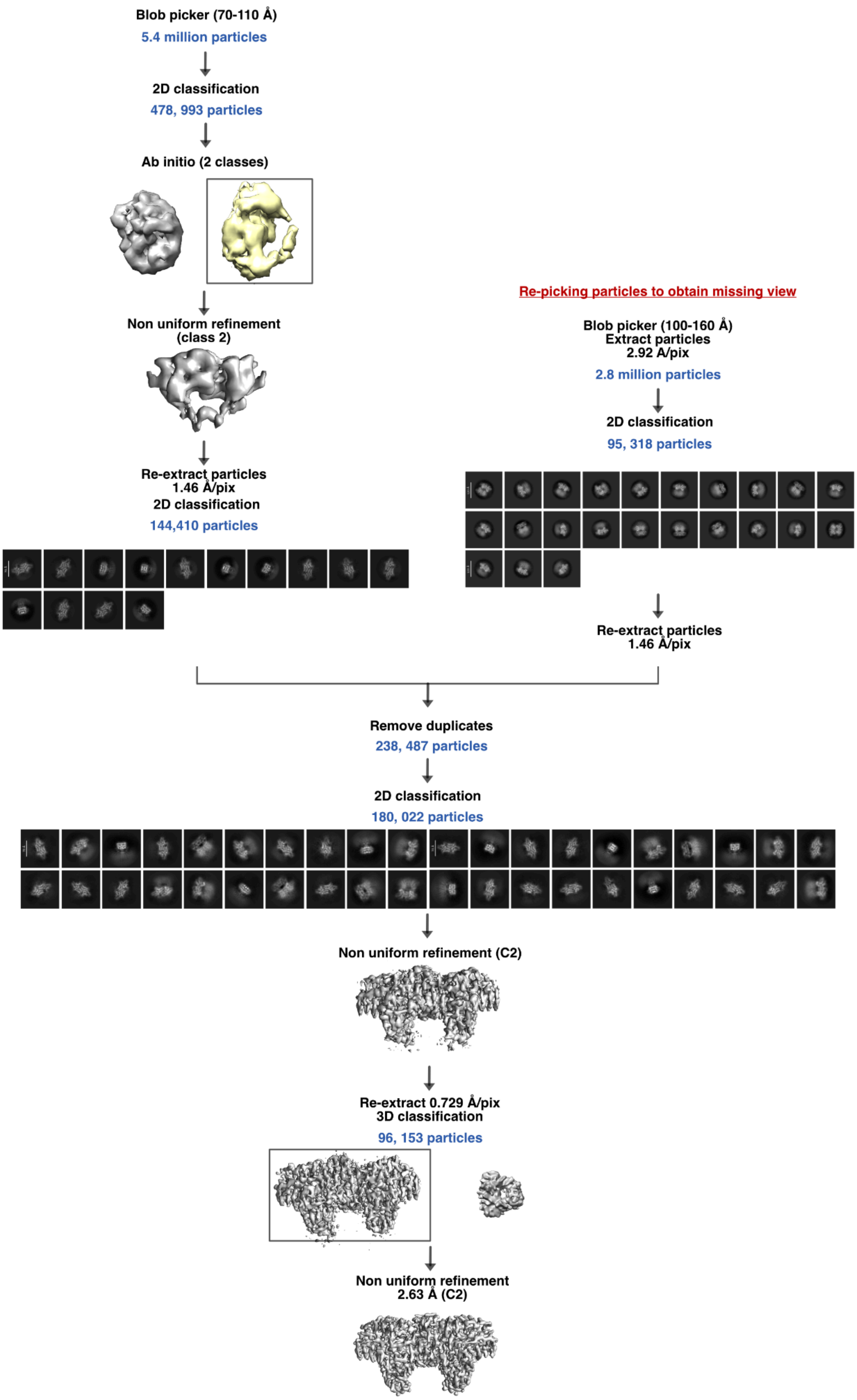


**Figure S2: Picking workflow for E320K PpfC$^{Cs}$.** Particle picking and classification was performed in CryoSPARC 4.5.3. The workflow is described in detail in the Materials and Methods section of the main paper.

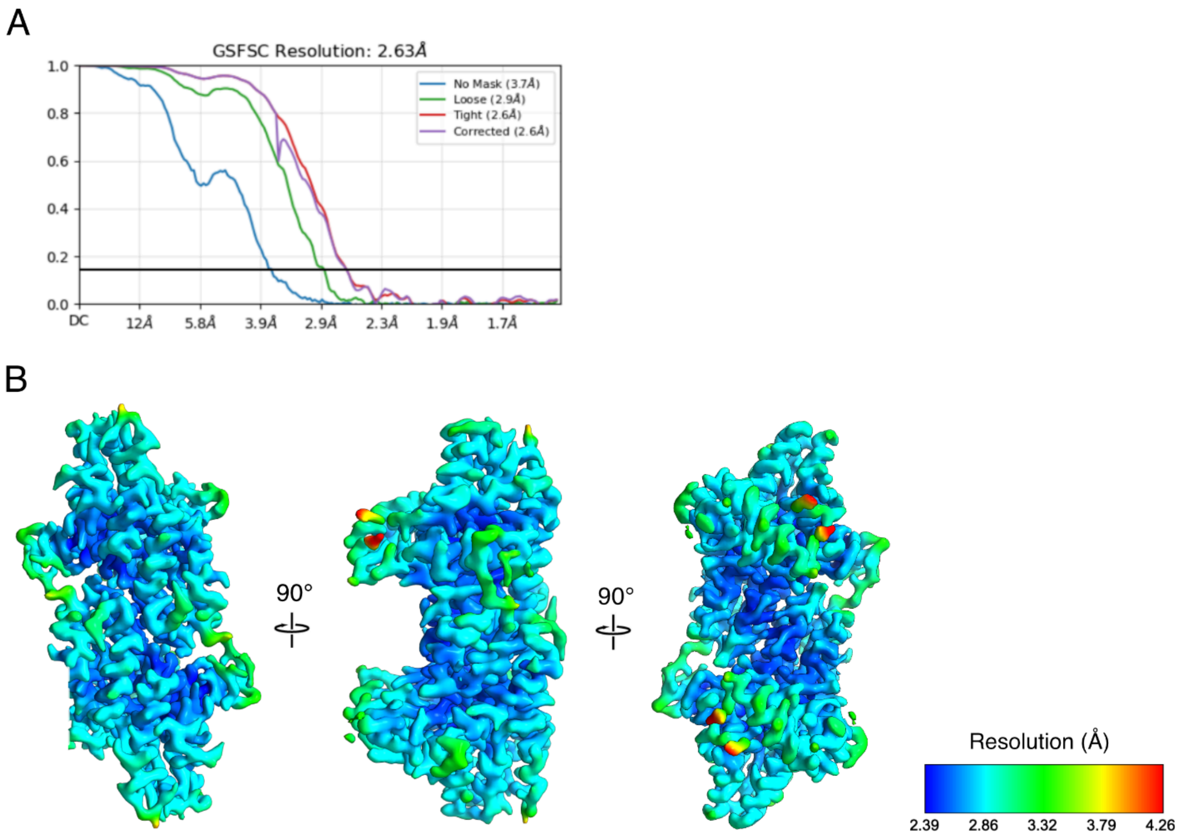


**Figure S3: FSC and local resolution of E320K PpfC$^{Cs}$. (*A*)** Gold standard fourier shell correlation for the E320K PpfC$^{Cs}$ dimer with C2 symmetry applied. The resolution was determined with FSC (Fourier Shell Correlation) = 0.143. A tight mask was required for refinement due to the proximity of particles to one another. **(*B*)** Local resolution was determined in PHENIX using the final unsharpened map. The resolution is shown using a rainbow palette, as indicated.

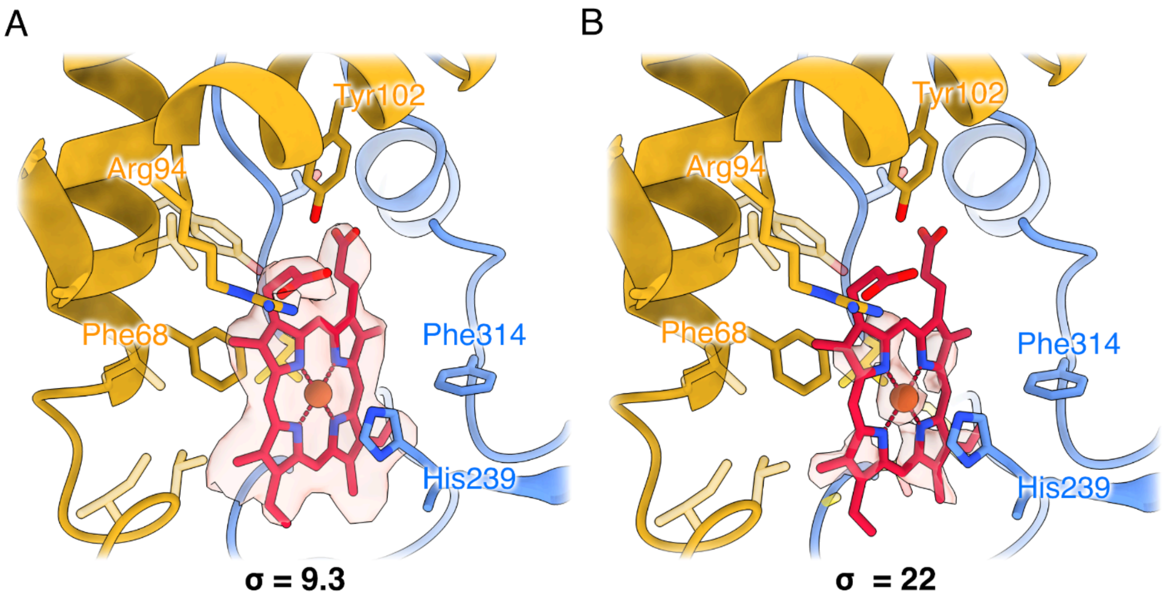


**Figure S4: Density for metalated PPIX at the active site of E320K PpfC$^{Cs}$.** View of the active site porphyrin (red with the central metal atom in orange), with the map contoured at two sigma values. The map at two different levels **(*A*, *B*)** clearly reveals density within the macrocycle, which we predict to be iron.

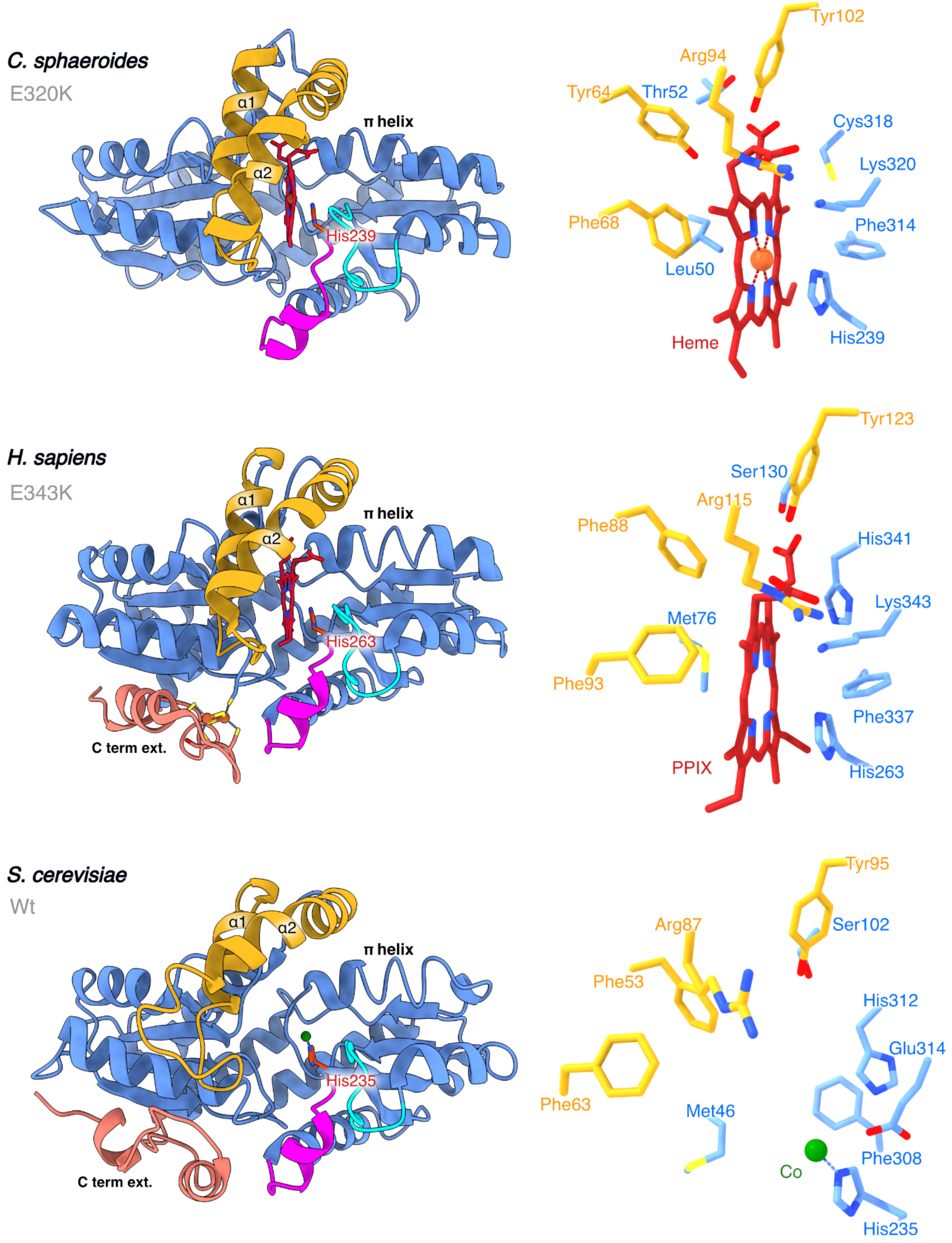


**Figure S5: Conserved PpfC architecture in PpfC$^{Cs}$, PpfC$^{Hs}$ and *Saccharomyces cerevisiae* PpfC (PpfC$^{Sc}$).** PpfC subunit architecture shown for E320K PpfC$^{Cs}$, E343K PpfC$^{Hs}$ (PDB: 1HRE) (19) and wild-type PpfC$^{Sc}$ (PDB: 1L8X) (17). A symmetric, bilobal domain surrounds the active site cleft, which is protected from bulk solvent by the highlighted upper and lower lip regions. As is described in the main text, the three lips are colored as follows: UL - gold, $LL_1$ - magenta, $LL_2$ - cyan. Active site residues for each variant are shown on the right. The π helix (35, 36) is shown to be in the wound conformation in both E320K PpfC$^{Cs}$ and E343K PpfC$^{Hs}$.

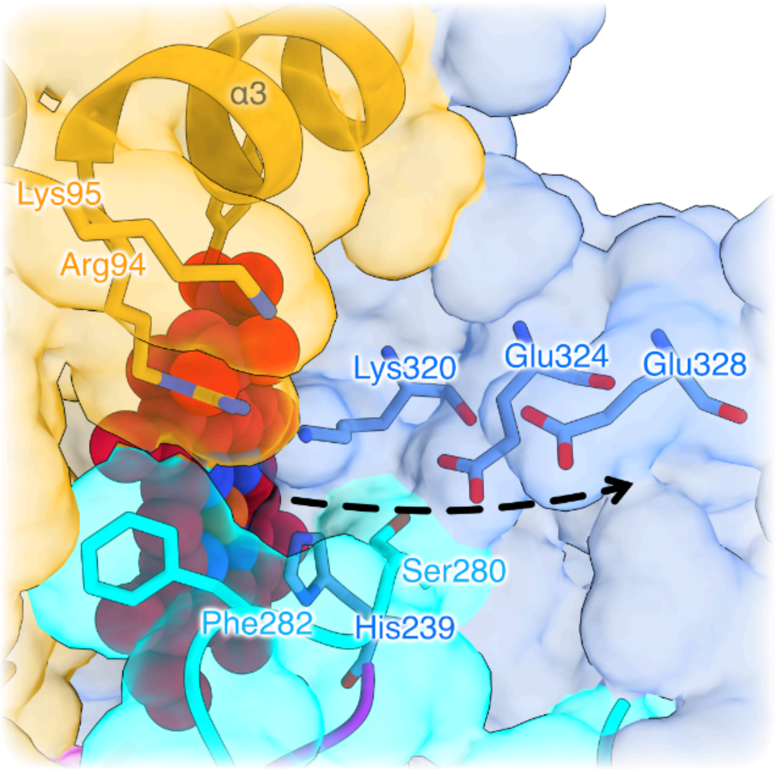


**Figure S6: Putative proton exit channel from His239 to the solvent.** Note that in the wild-type enzyme residue 320 is Glu not Lys. The arrow represents the proposed direction of proton exit, via Glu324 and Glu328. The enzyme is coloured as in Figure 2 of the main text.

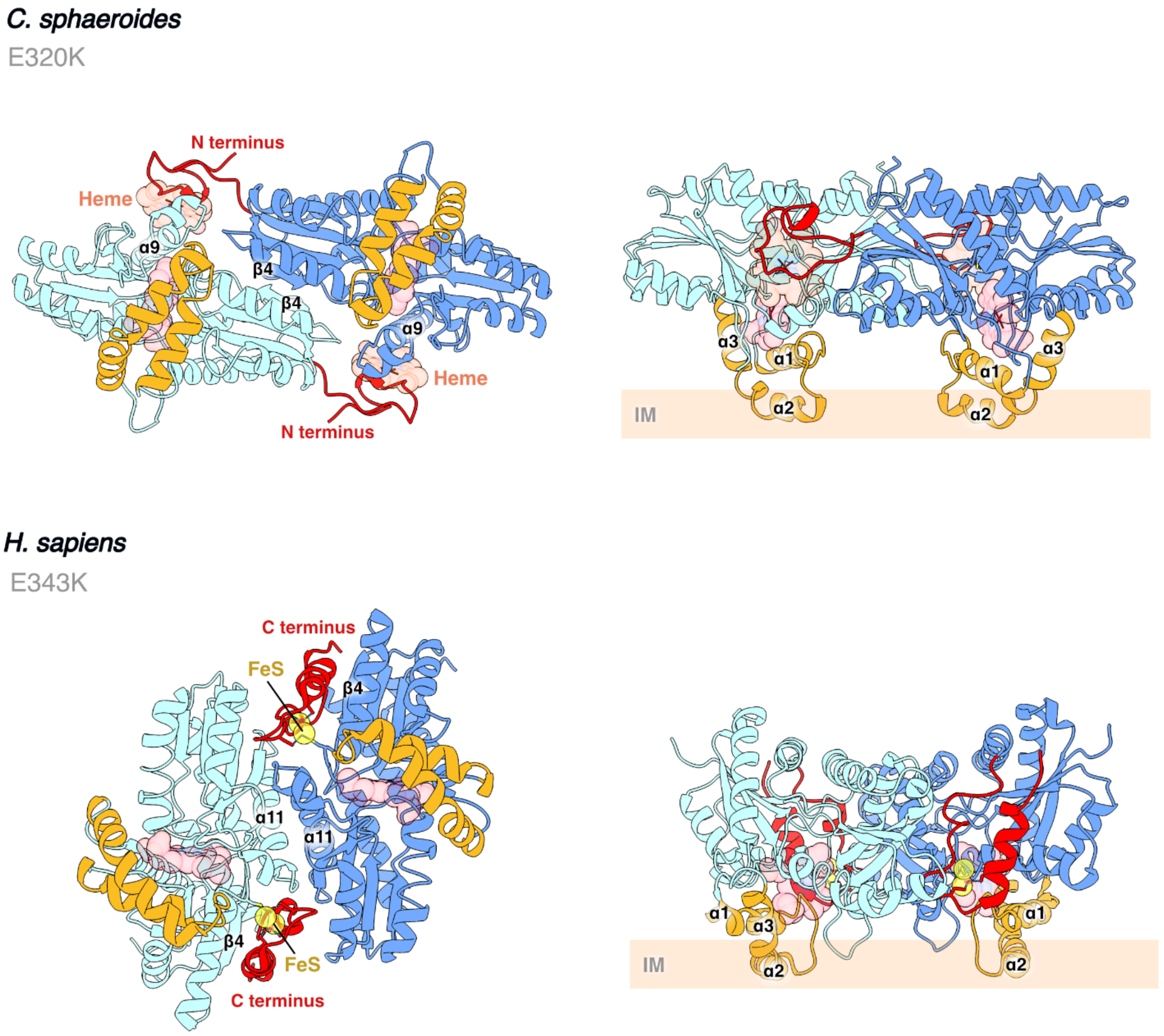


**Figure S7: Dimer interface comparisons between PpfC$^{Cs}$ and PpfC$^{Hs}$.** Cartoon representations of the E320K PpfC$^{Cs}$ and the E343K closed-state model of PpfC$^{Hs}$ (PDB:2HRE) (19). The respective N/C-terminal extensions for each species are shown in red. The presence of the C terminal extension of PpfC$^{Hs}$ precludes dimerization via the β4 sheets. Similarly, the N terminal extension of PpfC$^{Cs}$ precludes dimerization via the α9 helices (α11 in PpfC$^{Hs}$). Despite these differences, each dimer configuration allows for both active site lips (gold) to interface with the inner membrane (IM) in concert, which we propose (as has been suggested for PpfC$^{Hs}$ (19, 36)) allows hydrophobic substrate acquisition from, and product release to, the membrane environment.

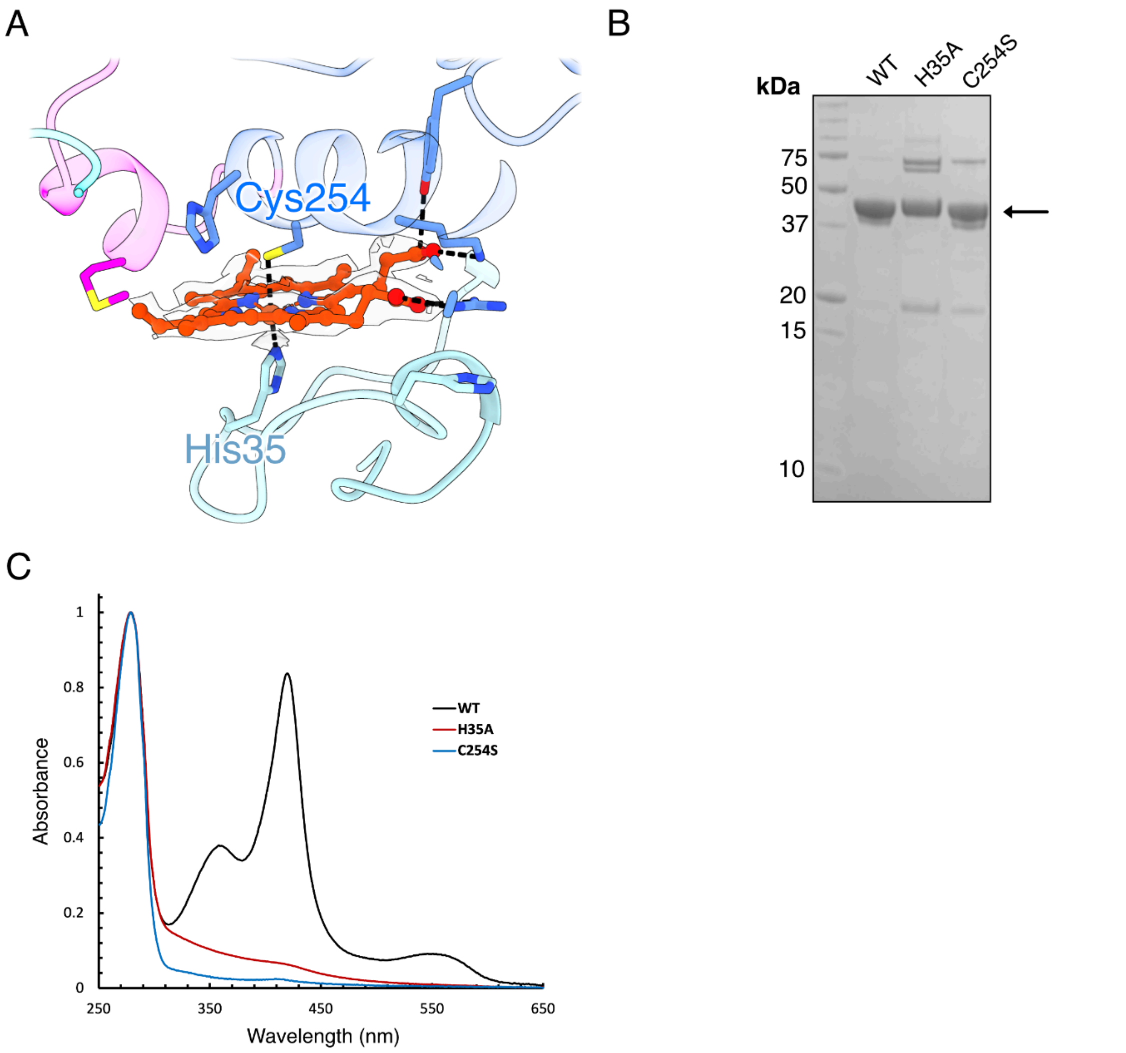


**Figure S8: Site directed mutagenesis of residues that coordinate the intra-dimer heme. (*A*)** View of the intra-dimer heme, showing coordination of the central iron by His35 (N terminal loop of one monomer) and Cys254 (α9 helix of the other monomer). **(*B*)** SDS-PAGE gel confirming similar purity of the three enzyme variants. **(*C*)** Absorbance spectra of isolated WT (black trace), H35A (red trace) and C254S (blue trace) variants of PpfC$^{Cs}$ showing the lack of heme binding in the two site-directed mutants. The spectra are normalized at 280 nm.

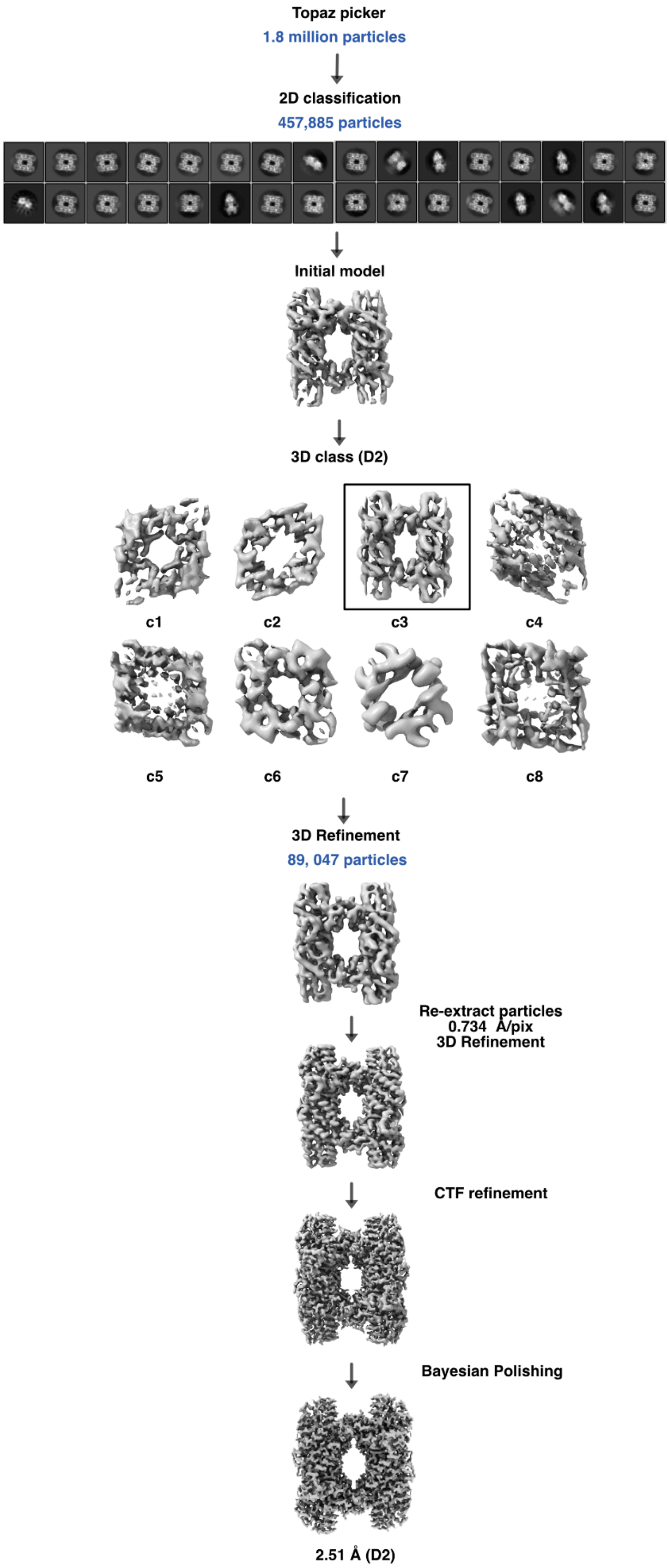


**Figure S9: Classification workflow for wild-type PpfC.** The PpfC tetramer was determined using RELION 4.1. The workflow is described in detail in the Materials and Methods section of the main paper.

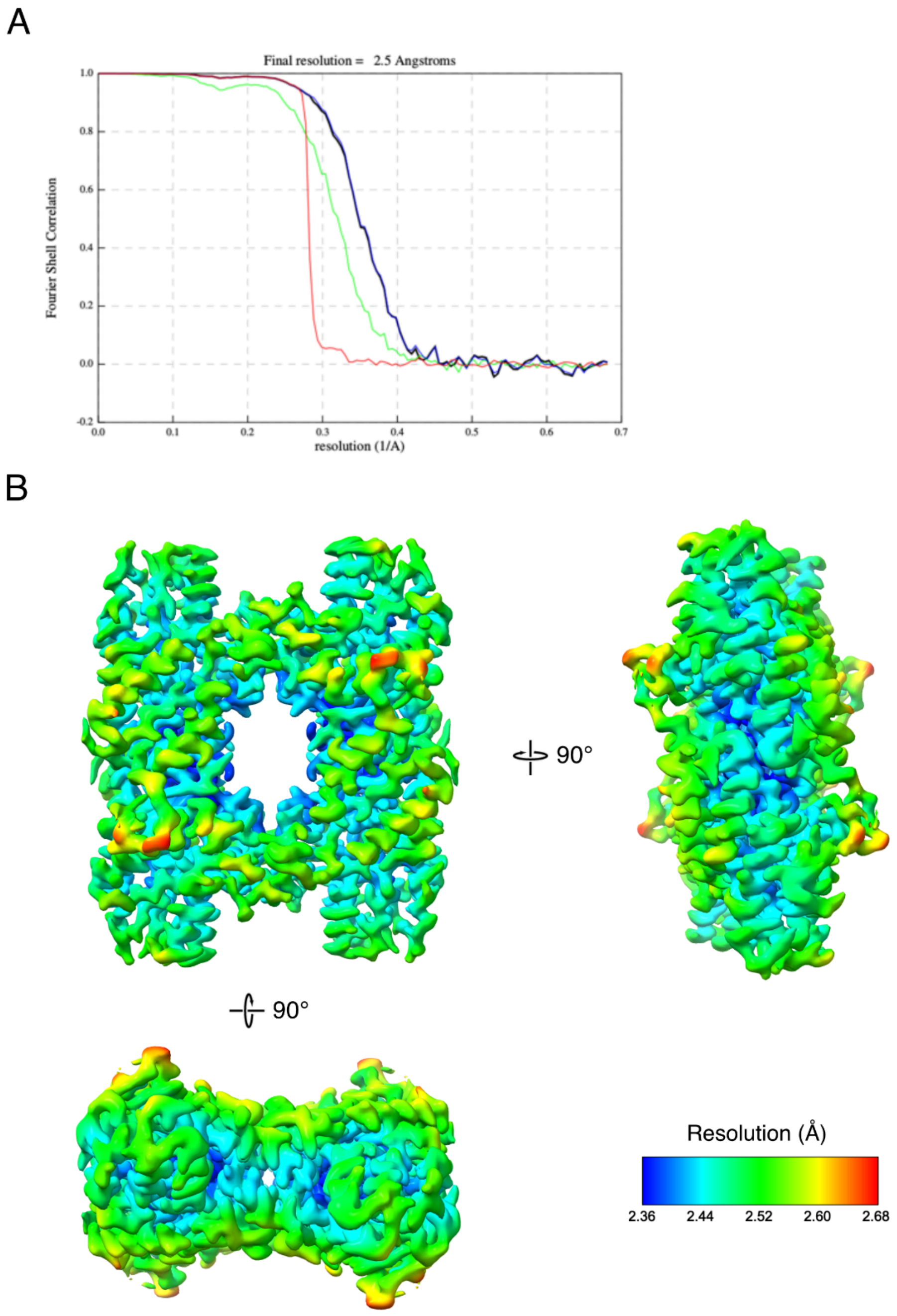


**Figure S10: FSC curve and local resolution estimates for the wild-type PpfC$^{Cs}$ map. (*A*)** The resolution was determined using RELION using the FSC (Fourier Shell Correlation) = 0.143. FSC curves are shown for unmasked (green), masked (black), phase-randomised (red), and mask-corrected (blue). **(*B*)** Local resolution was determined in PHENIX using the final unsharpened map. The resolution is shown using a rainbow palette. The resolution is shown using a rainbow palette, as indicated.

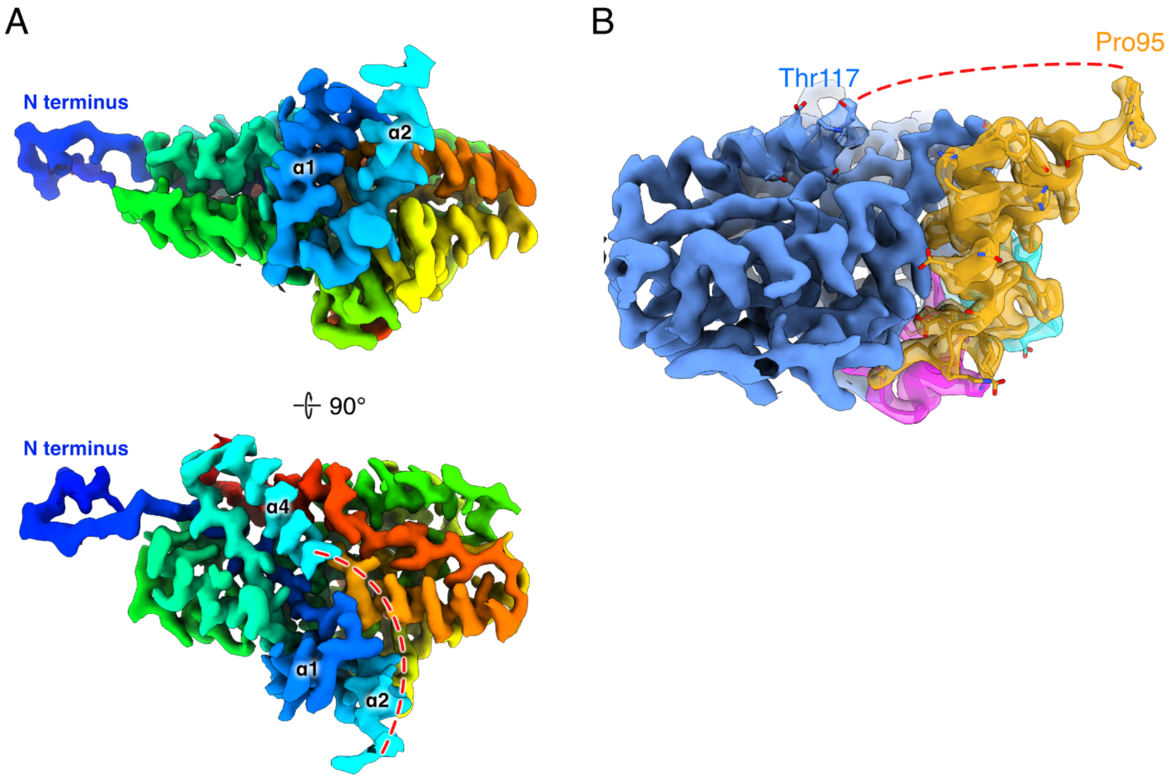


**Figure S11: Missing density in the upper lip region of wild-type PpfC$^{Cs}$. (*A*)** Cryo-EM density of a single monomer of wild-type PpfC$^{Cs}$, coloured by rainbow from blue (N terminus) to red (C terminus). The red dashed line indicates the region of missing density (residues 96-116). **(*B*)** View of the fitted model in the wild-type PpfC$^{Cs}$ cryo-EM density map, revealing the absence of density between Pro95 and Thr117.

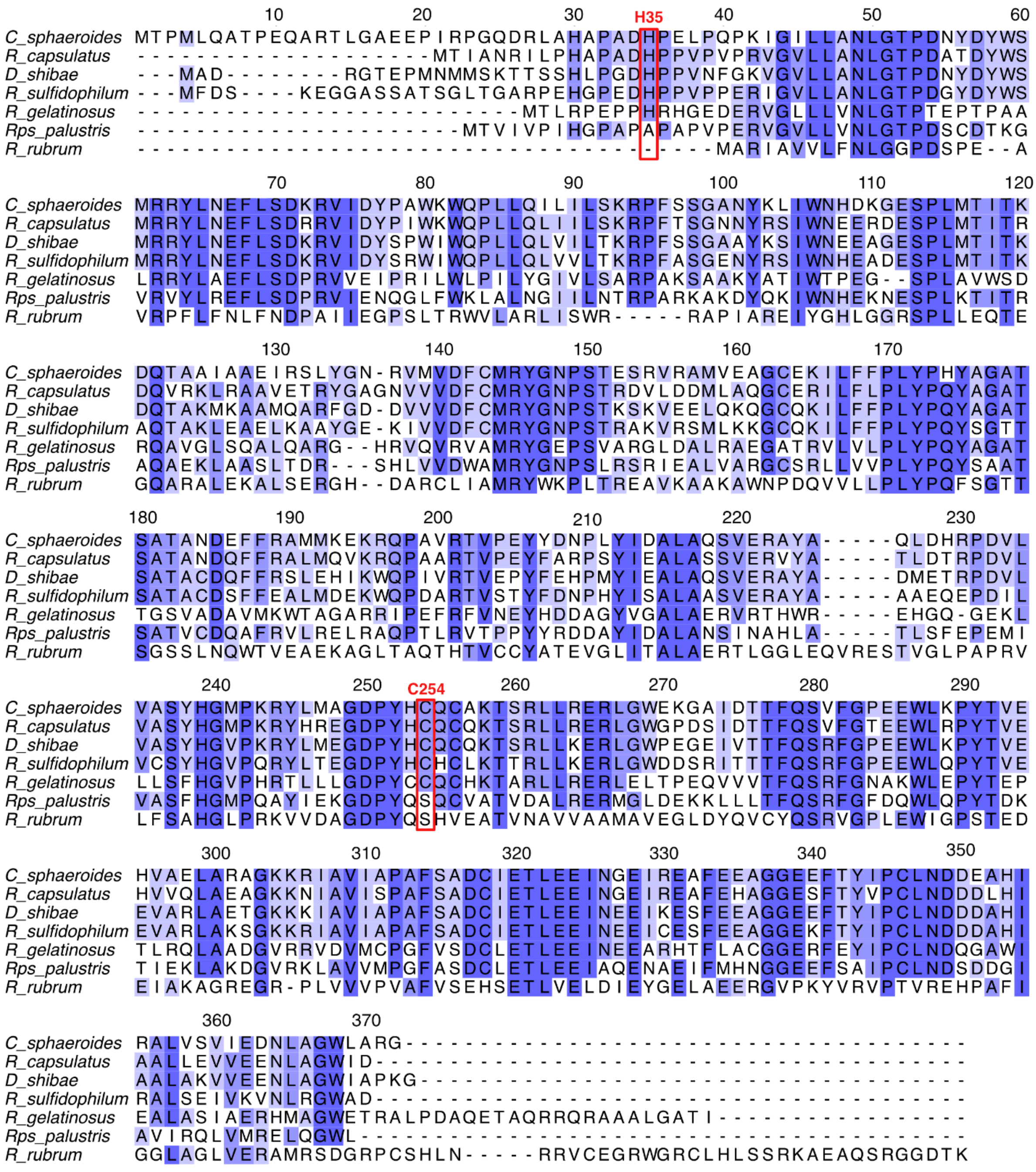


**Figure S12: Alignment of PpfC from selected purple bacteria.** PpfC from a range of purple bacteria were aligned using the Clustal Omega tool (53) and formatted in Jalview (54). Numbering represents the residue number of the *C. sphaeroides* PpfC. Aside from *R. rubrum* and *R. palustris*, the heme coordinating residues (His35 and Cys254 in $PpfC^{Cs}$; highlighted in red box) were well conserved in these examples.

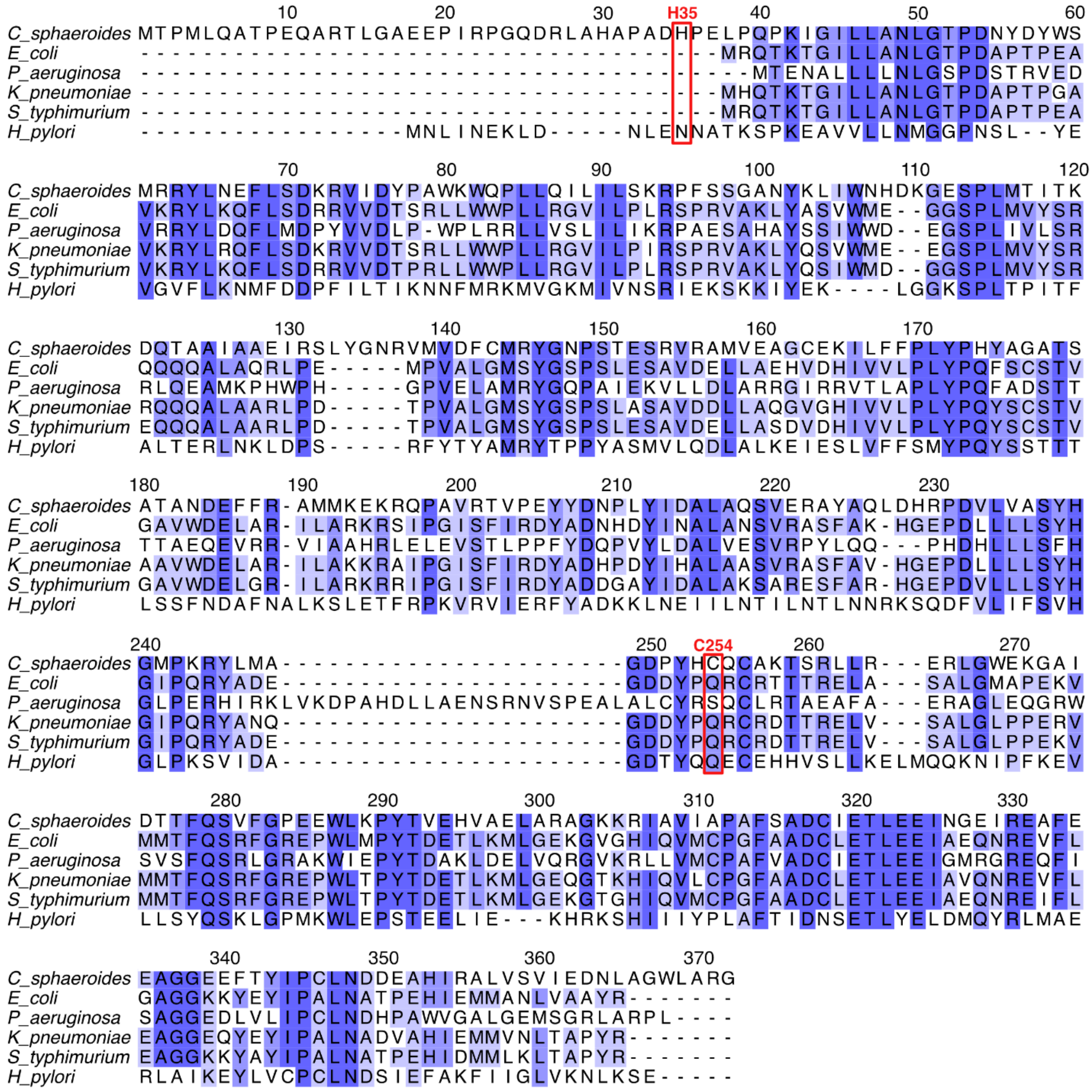

**Figure S13: Alignment of PpfC from selected Gram-negative bacteria.** PpfC from a range of Gram-negative bacteria were aligned using the Clustal Omega tool (53) and formatted in Jalview (54). Numbering represents the residue number of the *C. sphaeroides* PpfC. The heme coordinating residues (His35 and Cys254 in PpfC$^{Cs}$; highlighted in red box) were not conserved across the Gram-negative bacteria selected and appear to be a unique feature of some purple phototrophic bacteria.
